# Sequential eviction of crowded nucleoprotein complexes by the RecBCD molecular motor

**DOI:** 10.1101/142224

**Authors:** Tsuyoshi Terakawa, Sy Redding, Timothy D. Silverstein, Eric C. Greene

## Abstract

In physiological settings, all nucleic acids motor proteins must travel along substrates that are crowded with other proteins. However, the physical basis for how motor proteins behave in these highly crowded environments remains unknown. Here we use real–time single molecule imaging, kinetic Monte Carlo simulations, and Molecular dynamics simulations to determine how the ATP-dependent translocase RecBCD travels along DNA occupied by tandem arrays of high affinity DNA–binding proteins. We demonstrate that RecBCD forces each protein into its nearest adjacent neighbor, causing rapid disruption of the underlying protein–nucleic acid interface. This mechanism is not simply the same way that RecBCD disrupts isolated nucleoprotein complexes on otherwise naked DNA. Instead, molecular crowding itself completely alters the mechanism by which RecBCD removes tightly bound protein obstacles from DNA.

**Significance statement:** Chromosomes are crowded places, and any nucleic acid motor proteins that act upon DNA must function within these crowded environments. How crowded environments affect motor protein behaviors remains largely unexplored. Here, we use single molecule fluorescence microscopy visualize the ATP-dependent motor protein RecBCD as it travels along crowded DNA molecules bearing long tandem arrays of DNA-binding proteins. Our findings show that RecBCD can push through highly crowded protein arrays while evicting the proteins from DNA. Molecular dynamics simulations suggest that RecBCD forces the proteins into once another, causing rapid disruption of the protein-DNA interface. These findings may provide insights into how other types of motor proteins travel along crowded nucleic acids.

## Introduction

Long stretches of naked DNA do not exist in living cells. Instead, chromosomes are bound by all of the proteins that are necessary for genome compaction, organization, regulation and maintenance. DNA polymerases, RNA polymerases, helicases, translocases, chromatin-remodeling complexes, must all travel along the highly crowded nucleic acids that exist within these physiological settings. There is a growing appreciation that ATP-dependent motor proteins are required to either remove or remodel nucleoprotein complexes that may otherwise block normal processes related to nucleic acid metabolism, including DNA replication, transcription and DNA repair (1–9). Despite this importance, there remains almost no detailed mechanistic information describing how molecular motor proteins of any type behave on highly crowded nucleic acids.

RecBCD is a large (330–kDa) heterotrimeric complex that has served as an important model system for understanding the properties of nucleic acid motor proteins (10–13). RecBCD processes double–stranded DNA breaks (DSBs) during homologous recombination and replication fork rescue in *Escherichia coli*, and also degrades linear chromosome fragments to prevent aberrant DNA replication or recombination (10, 11, 14). Interestingly, RecB and RecC are the only two recombination proteins necessary for cell viability when head-on replication-transcription collisions are exacerbated by inversion of the ribosomal RNA operon (15). In addition to its roles in protecting genome integrity, RecBCD is also a self–defense enzyme that degrades foreign invaders, such as bacteriophage, and the resulting DNA fragments are incorporated into the CRIPSR locus, providing immunity against further infection (16). RecBCD possesses two ATP-dependent Superfamily 1 (SF1) molecular motor proteins, the 3’ →5’ SF1A helicase RecB (134-kDa), and the 5’→3 SF1B helicase RecD (67-kDa) (13). The RecC subunit (129-kDa) holds the complex together and coordinates the response to 8-nucleotide cis-acting Chi (crossover hotspot instigator) sequences (5’-dGCTGGTGG-3’). RecD is the lead motor before Chi, RecB is the lead motor after Chi and Chi recognition is accompanied by a reduced rate of translocation corresponding to the slower velocity of RecB (17, 18). RecB also contains a nuclease domain necessary for DNA-end processing, and recognition of Chi results in the production of 3’ single-stranded DNA overhangs onto which the recombinase RecA is loaded (10, 14).

All biological functions of RecBCD require it to travel along DNA that will be occupied by other proteins, and as such RecBCD has been used as a model for studying how motor proteins respond to DNA–bound obstacles (19, 20). Single–molecule observations have revealed that RecBCD can disrupt a variety of tenaciously bound nucleoprotein complexes, including EcoRI^E111Q^, RNA polymerase (RNAP), and lac repressor. RecBCD does not slow or pause during collisions with individual proteins. Instead, RecBCD appears to evict each of these different proteins through a common mechanism in which probability of protein dissociation is directly proportional to the number of steps they are forced to take as RecBCD pushes them from one nonspecific site to the next (19). RecBCD is even capable of stripping nucleosomes from DNA, highlighting it as an extremely powerful molecular motor (19, 20).

Here we sought to establish whether RecBCD could translocate along DNA substrates that were occupied by the high-affinity DNA-binding proteins EcoRI^E111Q^ or *E. coli* RNA polymerase holoenzyme. We demonstrate that under crowded conditions, RecBCD quickly and sequentially clears these nucleoprotein complexes from DNA by pushing adjacent proteins into one another. Our results suggest a model in which RecBCD uses DNA-bound proteins as molecular levers to generate torque, which rapidly destabilizes the underlying protein-nucleic acid interfaces. This unique mechanism of sequential protein disruption from the tandem arrays is entirely distinct from how RecBCD removes isolated protein complexes from DNA, indicating that molecular crowding itself can alter the mechanism by which ATP-dependent molecular motor proteins respond to nucleoprotein obstacles. These findings provide new insights into how molecular motors behave while traveling along nucleic acids in crowded physiological settings.

## Results

### Models for protein eviction in crowded environments

As an initial step towards understanding how motor proteins might behave in crowded settings, we first considered three generalized scenarios describing potential outcomes of RecBCD collisions with protein arrays (Fig. 1A). In the (*i*) accumulation model, RecBCD pushes each protein into its nearest neighbor without dislodging any of the proteins from the DNA, resulting in greater resistance as proteins continue to accumulate in front of the translocase. For the (*ii*) sequential model, RecBCD actively evicts each protein as it is encountered. In the (*iii*) spontaneous model, the proteins spontaneously dissociate according to their intrinsic dissociation rate constants, and RecBCD must wait for these dissociation events before moving forward. We include the spontaneous model as a formal possibility, although we note that this model is unlikely to be correct for RecBCD because previous experiments have shown that RecBCD can quickly push EcoRI^E111Q^, RNAP and lac repressor off of their respective cognate binding sites with no evidence of either slowing or pausing (19). Importantly, the accumulation and sequential models are not mutually exclusive. Indeed, RecBCD readily pushes isolated proteins for extended distances along DNA (19), suggesting that it might also push proteins into one another on crowded DNA. These observations suggest that some variation of the accumulation model could apply for RecBCD acting in crowded environments. To account for this possibility we also considered an alternative variation of the sequential model, in which small numbers proteins can accumulate in front of RecBCD before the accrued resistance leads to sequential dissociation (*SI Appendix*, Fig. S1A).

**Fig. 1.**
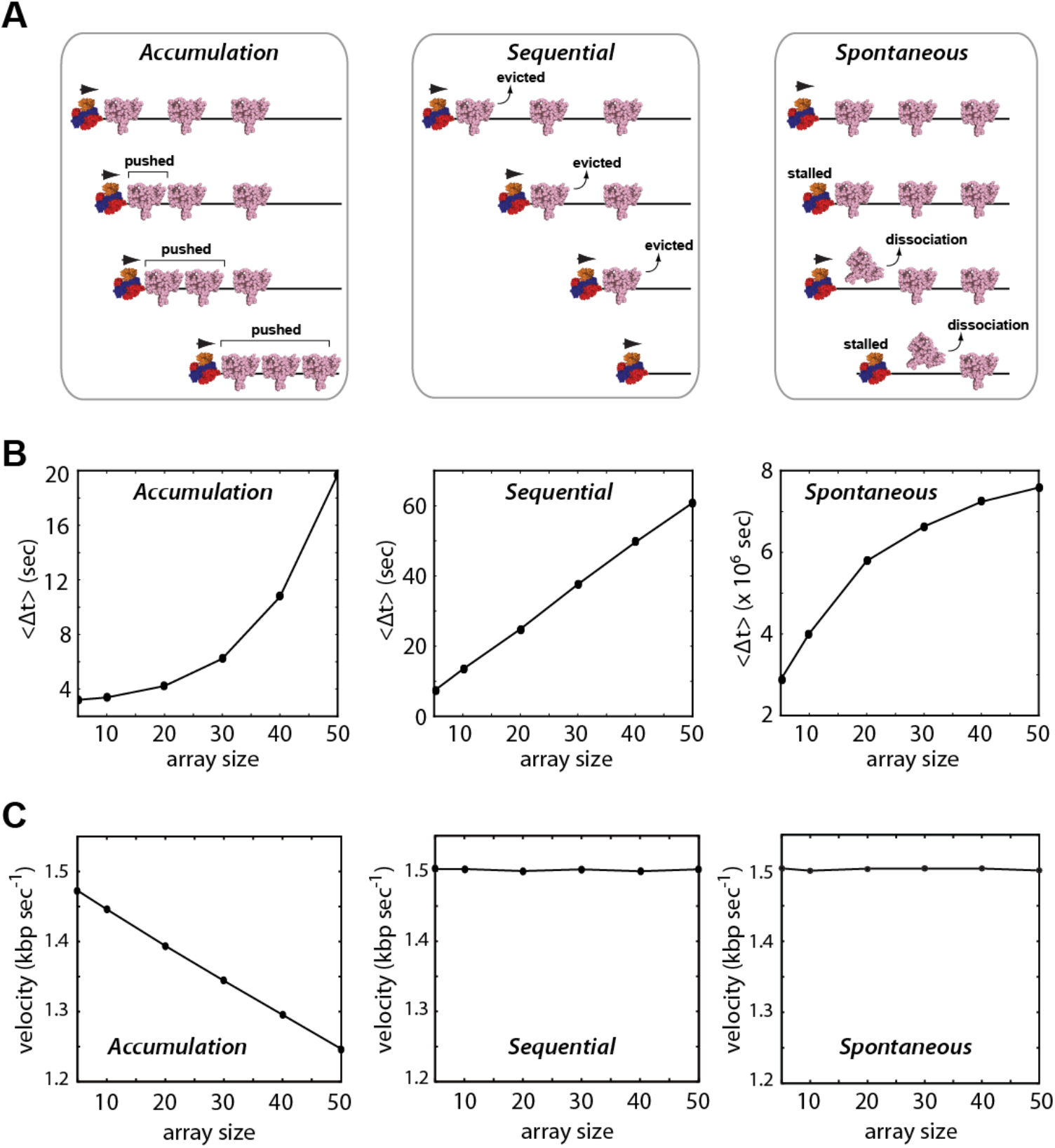
Models for translocase behavior in crowded environments. **(A)** Schematic depictions of three generalized models (accumulation, sequential, and spontaneous) for RecBCD movement through extend protein arrays. Details of each model are presented in the main text. **(B)** Results from kinetic Monte Carlo simulations for each different model showing the predicted relationships between RecBCD pause durations (〈Δt〉) and protein array size. **(C)** Predicted post-array RecBCD velocities for each model.

### Monte Carlo simulations of protein eviction by DNA translocases

We next performed Kinetic Monte Carlo (KMC) simulations as a means to predict potential experimental outcomes for each of the different models. For these KMC simulations, we modeled the behavior of RecBCD on DNA substrates bearing 5-50x tandem arrays of the high-affinity DNA-binding protein EcoRI^E111Q^. This protein is a catalytically inactive version of the EcoRI restriction endonuclease that binds tightly to DNA, but is unable to cleave its cognate target site (21). We chose EcoRI^E111Q^ for our computational and experimental studies because it is one of the highest affinity DNA–binding proteins known to exist, with site–specific and non–specific dissociation constants (*K*_d_) of ~2.5 femtomolar (fM) and ~4.8 picomolar (pM) (21), respectively. EcoRI^E111Q^ is also a highly potent block to both the transcription (22–24) and DNA replication machineries (25, 26). In addition, wild–type EcoRI can withstand up to ~20–40 pN of applied force (27). EcoRI^E111Q^ binds to its cognate target ~3000–fold more tightly than wild–type EcoRI, therefore, we infer that EcoRI^E111Q^ can resist at least as much force as the wild–type protein.

Within each KMC simulation, the DNA–bound proteins must either slide or dissociate upon collision with RecBCD. The accumulation model is realized by prohibiting dissociation of the obstacle proteins, which are instead always pushed by RecBCD. The sequential model requires RecBCD to provoke protein dissociation prior to moving forward. In the spontaneous model, RecBCD must wait until proteins dissociate according to their intrinsic dissociation rate constants. Each model predicts that RecBCD will slow or stall upon encountering the array; these events should be revealed as an experimentally observable pause coinciding with the location of protein array (*SI Appendix*, Fig. S1A). Importantly, the accumulation, sequential, and spontaneous dissociation models all yield distinct predictions for the relationship between apparent pause duration (〈Δt〉) and array size (Fig. 1B and *SI Appendix*, Fig. S2B). The accumulation model predicts an exponential increase in pause duration with increasingly large protein arrays, the sequential model predicts a linear relationship between pause duration and array size, and the spontaneous model predicts a logarithmic variation in pause duration for the different array sizes. The spontaneous dissociation model also predicts RecBCD will traverse the array orders of magnitude more slowly than the other models because RecBCD must wait for each protein to dissociate (Fig. 1B). In addition, the accumulation model predicts that RecBCD will experience a persistent reduction in velocity after traversing the array due to the accumulated resistance of the proteins it must push as it continues to move along the DNA (Fig. 1C). Finally, if small numbers of EcoRI^E111Q^ can build up in front of RecBCD prior to dissociating, then the pause duration will scale approximately linearly with array size as up to five proteins accumulate in front of RecBCD, similar to expectations for a purely sequential model (*SI Appendix*, Fig. S1B). However, pause duration begins to scale exponentially if larger numbers of proteins can accumulate in front of RecBCD before dissociation, yielding results that would be more similar to the pure accumulation model (*SI Appendix*, Fig. S1B).

### Visualizing removal of unlabeled EcoRI^E111Q^ from crowded DNA

We next sought to establish an experimental approach for directly testing the predictions for each of the different models. To accomplish this we engineered λ-phage DNA molecules bearing tandem arrays of 5, 10, 30 or 50 EcoRI binding sites. We then used single molecule DNA curtain assays to experimentally determine whether quantum dot (Qdot)–tagged RecBCD could traverse these protein arrays (Fig. 2A), and if so, determine which of the models presented above might most closely reflected the experimental data.

**Fig. 2.**
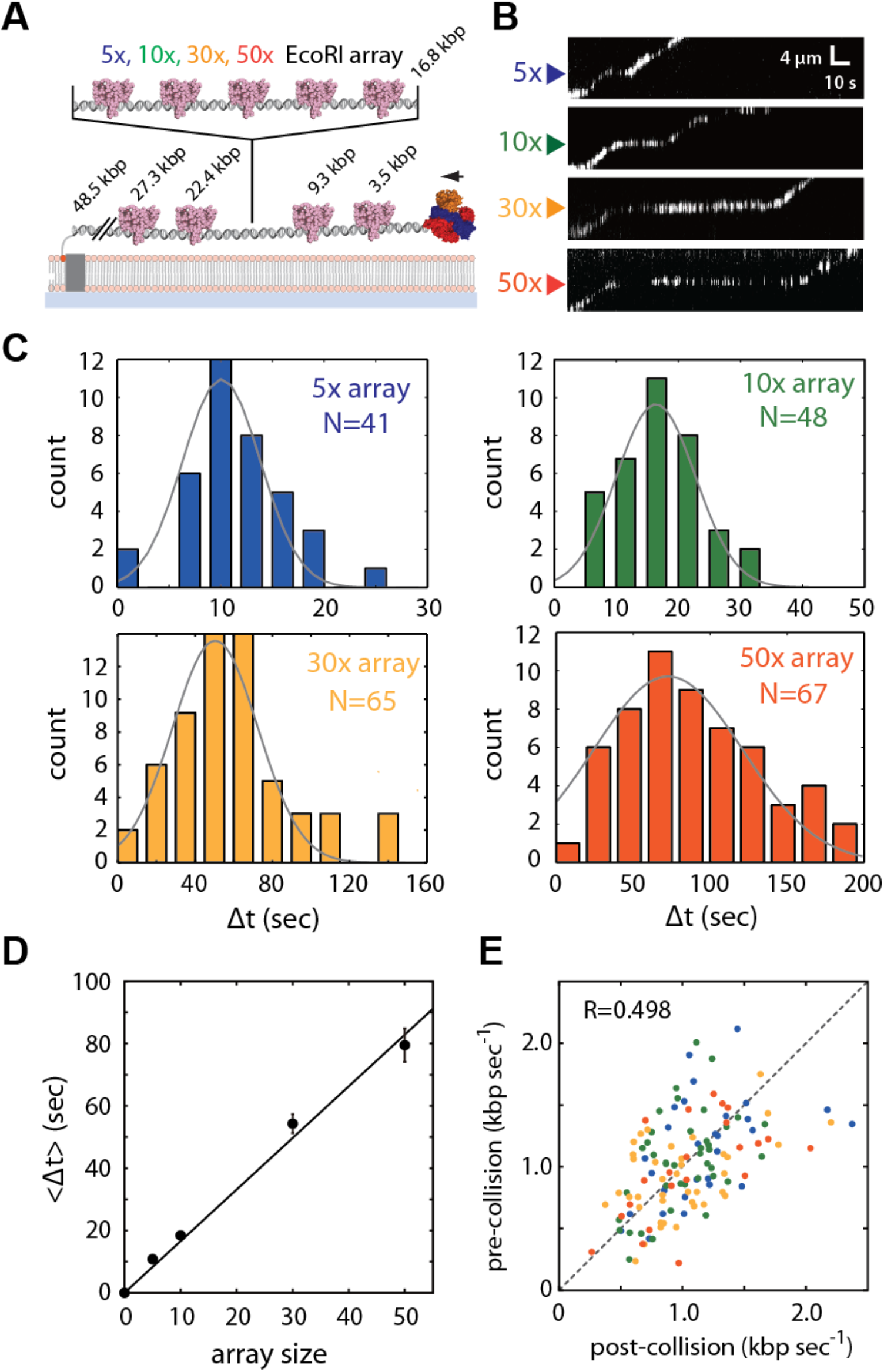
RecBCD can traverse highly crowed DNA substrates. **(A)** Schematic of the DNA curtain assay used to assess RecBCD behavior during passage through protein arrays. The DNA contains four native EcoRI binding sites (as indicated) and a cloned array of 5–50x EcoRI binding sites. The reactions were initiated by addition of 1 mM ATP into the RecBCD buffer (40 mM Tris–HCl [pH 7.5], 2 mM MgCl_2_, and 0.2 mg ml^−1^ Pluronic). **(B)** Representative kymographs showing RecBCD movement through 5–50x arrays in the presence of saturating EcoRI^E111Q^. Gaps in the RecBCD trajectories result from Qdot blinking. **(C)** Experimentally observed pause distributions for each array. Each data set is fitted by a Gaussian distribution to derive the average pause duration. **(D)** Experimental pause duration (〈Δt〉) plotted against array size. Error bars represent s.d. derived from a bootstrap analysis. **(E)** Scatter plot showing RecBCD pre– and post–collision velocities; color-coding is the same as in **(B)** and **(C)**.

In the absence of EcoRI^E111Q^, Qdot–tagged RecBCD displayed high processivity and monotonic translocation with two peaks in the velocity distribution, corresponding to 745±37 (13.6%) and 1,368±18 bp sec^−1^ (86.4%), in good agreement with reports for the properties of unlabeled RecBCD on naked DNA (*SI Appendix*, Fig. S3A and B) (18, 28, 29). We next asked whether the presence of EcoRI^E111Q^ affected the translocation behavior of RecBCD. Remarkably, RecBCD was still able to processively translocate along the DNA in the presence of saturating EcoRI^E111Q^ concentrations (Fig. 2B and *SI Appendix*, Fig. S4). However, as predicted by the KMC simulations, RecBCD exhibited a pronounced pause upon encountering each of the protein arrays (Fig. 2B and C), and the average pause duration (〈Δt〉) scaled linearly with array size (Fig. 2D). Control experiments with unlabeled RecBCD and unlabeled EcoRI^E111Q^ were in close agreement with results from Qdot-tagged RecBCD, arguing against the possibility that the Qdot might alter the outcomes of the collisions (*SI Appendix*, Fig. S5). In addition, RecBCD slowed to an average apparent velocity of just 27.3±3.4 bp sec^−1^ while traversing the EcoR^E111Q^ arrays, but resumed its pre–collision velocity once beyond the array. This finding indicates that encounters with the tandem high-affinity nucleoprotein complexes had no lasting impact on the translocation properties of the enzyme (Fig. 2E and *SI Appendix*, Fig. S3C). Together, these results are in closest agreement with expectations for the sequential eviction model, or a variation of the sequential model involving the accumulation of a small number of EcoRI^E111Q^ dimers in front of RecBCD (Fig. 1B and C and *SI Appendix*, Fig. S1B). We do not yet know precisely how many EcoRI^E111Q^ dimers might accumulate in front of RecBCD before they start dissociating. However, DNA curtain experiments have revealed that RecBCD can concurrently push at least two EcoRI^E111Q^ dimers (*SI Appendix*, Fig. S1C), whereas the tandem arrays of five EcoRI^E111Q^ dimers cause a pronounced pause in RecBCD translocation (Fig. 2). Therefore, we conclude that between two to five EcoRI^E111Q^ dimers might accumulate in front of RecBCD before the combined resistance leads to their sequential eviction.

### Sequential disruption of fluorescent EcoRI^E111Q^ arrays

The KMC simulations and experimental results presented above, together with the observation that RecBCD readily pushes isolated molecules of EcoRI^E111Q^ (19), all suggest that RecBCD sequentially removes proteins from crowded DNA, and that it may do so by pushing the proteins into one another. This model makes at least two important predictions that can be experimentally tested using DNA curtain assays. First, the sequential models predict that passage of RecBCD through a fluorescently tagged protein array should coincide with a linear decrease in the array signal as the proteins are evicted, but the last protein(s) within the array should be pushed for long distances along the naked DNA because they will encounter no resistance from more distal obstacles (19) (Fig. 3A). To test this first prediction, we labeled a λ-DNA substrate bearing a 50x EcoRI binding site array with Qdot -tagged EcoRI^E111Q^, and then asked whether and how unlabeled RecBCD passed through these arrays. As anticipated, eviction of the fluorescent proteins from the 50x array was initially observed as a linear decrease in the overall fluorescence signal intensity as unlabeled RecBCD moved slowly through the array (Fig. 3B). Also as predicted, upon reaching the end of the array, RecBCD resumed its normal velocity and pushed the remaining protein(s) towards the end of the DNA molecule (Fig. 3B).

**Fig. 3.**
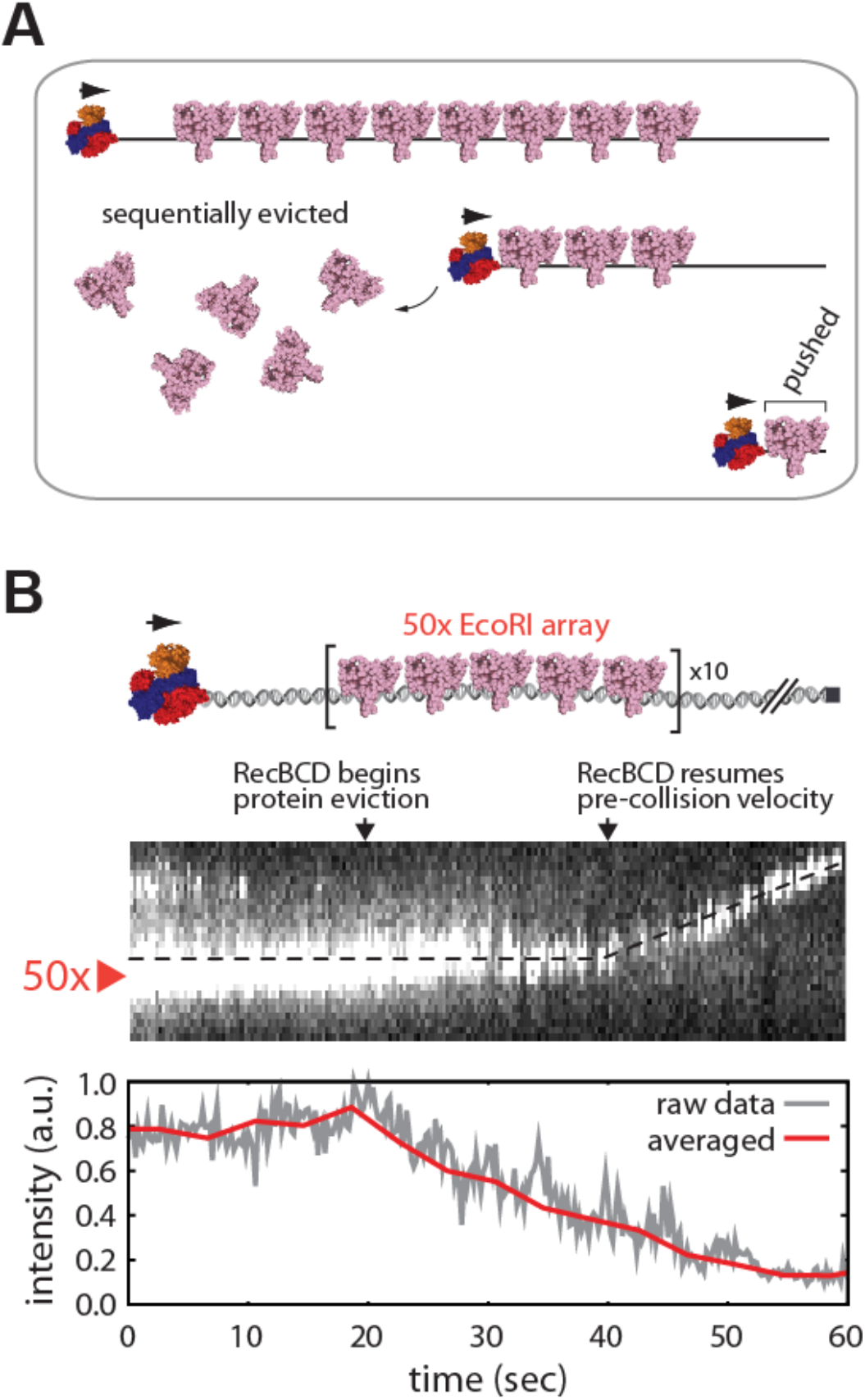
RecBCD collisions with fluorescent 50x arrays of EcoRI. **(A)** Model describing predicted outcome for RecBCD collisions with the 50x EcoRI^E111Q^ arrays; details are presented in the main text. **(B)** Representative kymograph and corresponding signal intensity profile showing unlabeled RecBCD traversing a 50x array of Qdot–tagged EcoRI^E111Q^. Reactions were initiated by addition of 1 mM ATP into the RecBCD buffer (40 mM Tris–HCl [pH 7.5], 2 mM MgCl_2_, plus 0.2 mg ml^−1^ Pluronic). Gaps in the EcoRI^E111Q^ trace result from Qdot blinking.

The sequential models also predicts that if RecBCD pushes a single proximal EcoRI^E111Q^ into a more distal EcoRI^E111Q^ array, then the resulting collision should coincide with preferential eviction of the proximal protein as it is driven into the larger array (Fig. 4A). To test this prediction, we performed two-color single molecule experiments in which separate aliquots of EcoRI^E111Q^ were labeled with either green (Qdot 605) or red (Qdot 705) quantum dots. The differentially labeled proteins were bound to λ-DNA bearing the 5x EcoRI binding site array. We then visually identified DNA molecules with appropriately dispersed mixtures of red and green EcoRI^E111Q^ bound to the native EcoRI sites within the λ-DNA and the engineered 5x arrays (Fig. 4B). As predicted by sequential models, when unlabeled RecBCD pushed Qdot–tagged EcoRI^E111Q^ into a 5x array, the proximal protein rapidly dissociated from the DNA upon encountering the 5x array in ~93% of experimentally observed collisions (N=25/27) (Fig. 4B). We also compared the fates of isolated EcoRI^E111Q^ molecules bound to the native EcoRI sites located either upstream or downstream of the engineered 5x array (Fig. 4C). As predicted by the sequential model, the upstream proteins most commonly dissociated from the DNA upon being pushed into the 5x EcoRI^E111Q^ arrays (Fig. 4C and D). In striking contrast, the downstream proteins were pushed for much longer distances and often survived until reaching the chromium (Cr) barrier at the tethered ends of the DNA molecules (Fig. 4C and D). Taken together, these experimental findings all support a model in which RecBCD clears crowded DNA of tightly bound EcoRI^E111Q^ by pushing the proteins into one another, resulting rapid and sequential removal of proteins from the DNA.

**Fig. 4.**
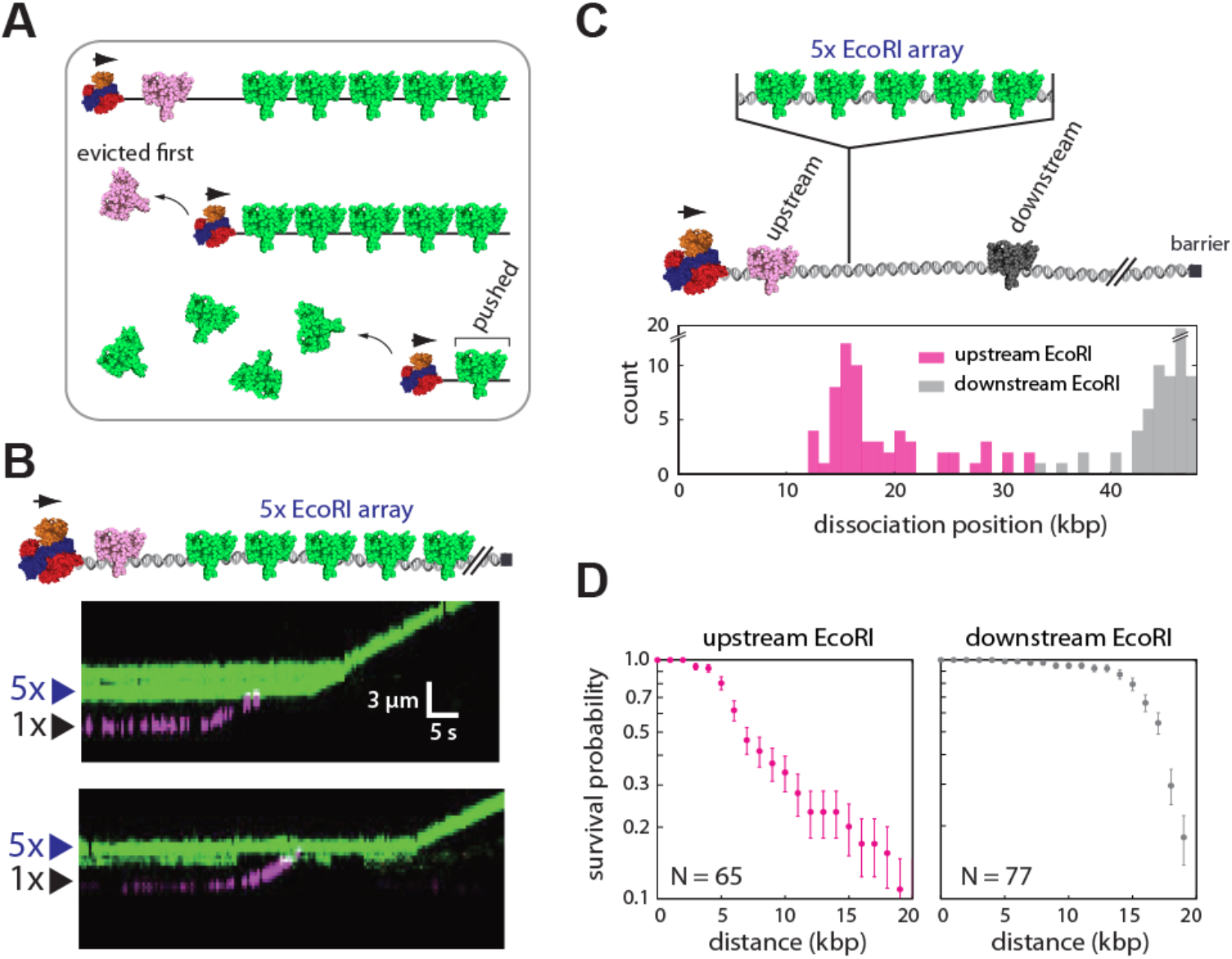
Sequential eviction of EcoRI from DNA by RecBCD. **(A)** Model describing the predicted outcome for two-color labeling experiments designed to test for sequential protein eviction by RecBCD; details are presented in the main text. **(B)** Examples of two–color kymographs showing Qdot–tagged EcoRI^E111Q^ (Qdot 705; magenta) being pushed into a 5x array bound by Qdot −tagged EcoRI^E111Q^ (Qdot 605; green). Reactions were initiated by addition of 1 mM ATP into the RecBCD buffer (40 mM Tris–HCl [pH 7.5], 2 mM MgCl_2_, plus 0.2 mg ml^−1^ Pluronic). Gaps in the EcoRI^E111Q^ traces result from Qdot blinking. **(C)** Schematic illustration of the experiment used to assess the fates EcoRI^E111Q^ bound to native target sites in the λ-DNA located either upstream or downstream of the 5x EcoRI array (upper panel), and resulting dissociation positions of the upstream and downstream EcoRI^E111Q^ molecules (lower panel). **(D)** Survival probability plots for EcoRI^E111Q^ molecules located upstream and downstream of the 5x EcoRI arrays.

### Molecular dynamics simulations of protein-protein collisions

The molecular events that take place during the collisions would be extremely difficult, if not impossible, to analyze experimentally due to the short spatial and temporal scales. Therefore, we used Molecular Dynamics (MD) simulations to more closely examine what might take place when RecBCD encounters with EcoRI^E111Q^ in crowded settings. The MD simulations utilized coarse–grained representations of the DNA and EcoRI^E111Q^, and collisions were conducted on 100-or 150-bp DNA fragments bearing either two or three EcoRI binding sites, respectively (Fig. 5). The interaction parameters for the proximal EcoRI^E111Q^ molecule(s) recapitulated the experimentally observed specific and non–specific EcoRI^E111Q^ dissociation constants (*SI Appendix*, Fig. S6), and the distal EcoRI^E111Q^ was bound more tightly to the DNA to emulate the accumulated resistance of several nucleoprotein complexes. RecBCD was modeled as a simple torus placed at one end of DNA, and exerted a constant force of 100 pN as it pushed against the DNA–bound molecules of EcoRI^E111Q^. The force that RecBCD can exert upon colliding with a DNA-bound obstacle is unknown, so we chose 100 pN for the MD simulations because this value is within an order of magnitude of several of the most powerful molecular motors that have been characterized (30). The expectation that RecBCD would be capable of exerting a force comparable to the most powerful molecular motor proteins is consistent with the finding that RecBCD can clear DNA bound by 50x arrays of EcoRI^E111Q^, and is also consistent with the finding that RecBCD readily overwhelms the hexameric motor protein FtsK, which itself exhibits a stall force of ~65 pN (31).

**Fig. 5.**
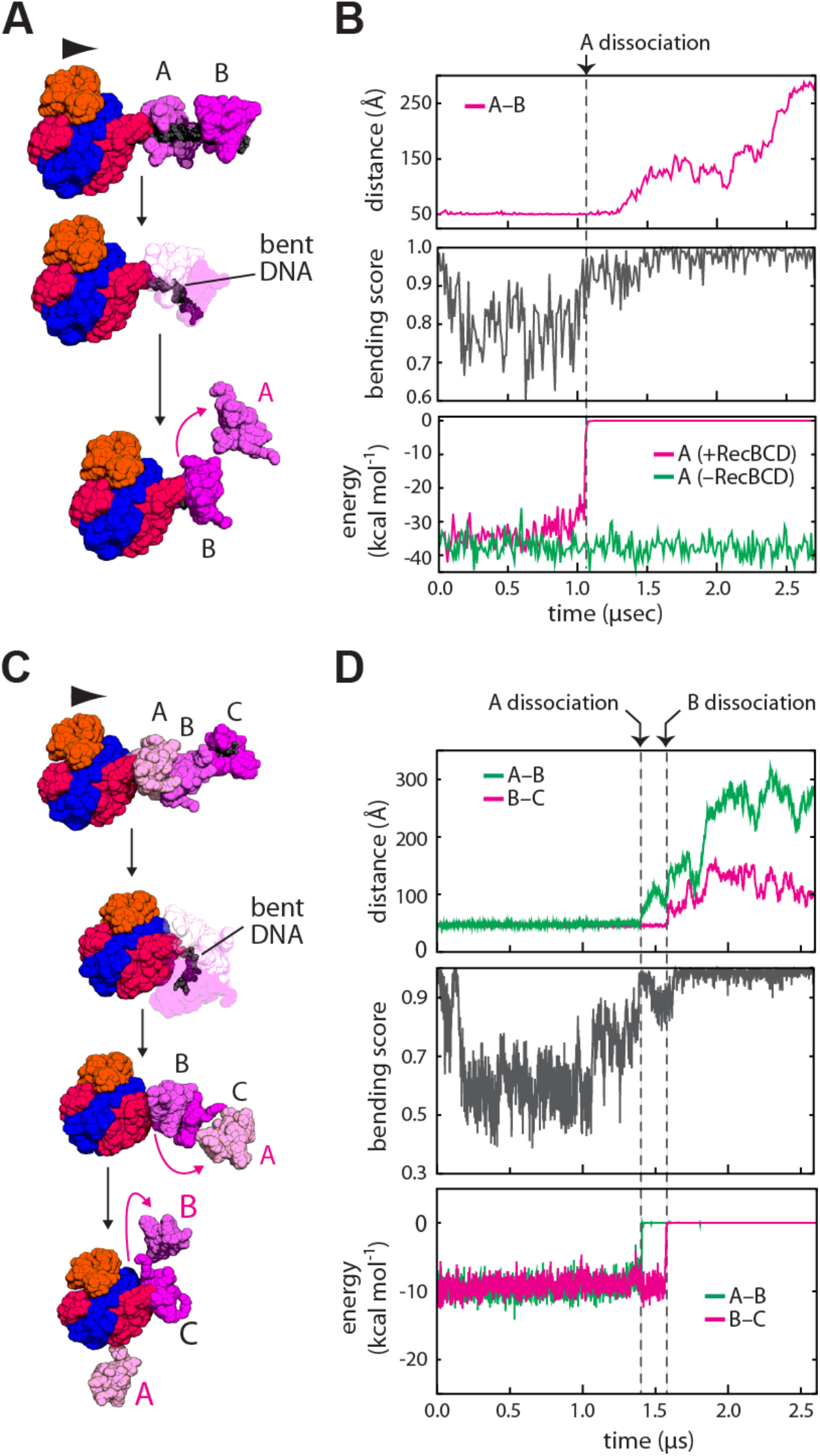
Molecular dynamics simulations of protein-protein collisions. **(A)** Snapshots from a representative simulation trajectory involving two EcoRI^E111Q^ molecules (also see *SI Appendix*). RecBCD begins at one end of the DNA and an arrowhead indicates its direction of travel. The proximal and distal proteins are labeled A and B, respectively. Magenta arrowheads and lettering highlight protein eviction. **(B)** The time trajectories showing the distance between the two EcoRI^E111Q^ molecules (top), the DNA bending score (middle), and interaction energy between the proximal EcoRI^E111Q^ and DNA (bottom). The zero time point corresponds to the initial collision of proteins A and B. **(C)** Snapshots from a representative simulation trajectory involving three EcoRI^E111Q^ molecules, labeled A, B and C, as indicated. (also see *SI Appendix*). **(D)** Time trajectories showing the distance between the EcoRI^E111Q^ molecules (top; A–B distance, green line; B–C distance, magenta line), DNA bending score (middle), and interaction energy between the EcoRI^E111Q^ molecules (A, green line; B, magenta line) and DNA (bottom). The zero time point corresponds to the initial collision of proteins A, B and C.

A representative MD simulation trajectory of RecBCD traveling on DNA bound by two EcoRI^E111Q^ molecules is shown in Fig. 5A and *SI Appendix*. As anticipated, RecBCD pushed the proximal EcoRI^E111Q^ along the DNA major groove until it collided with the distal EcoRI^E111Q^. The proximal protein then dissociated from the DNA as it was forced against the distal EcoRI^E111Q^ in all simulations (N=15/15) (Fig. 5B). The simulation time scale for each dissociation event was on the order of ~1.0 μsec (Fig. 5B), corresponding to ~10^8^–fold rate enhancement relative to spontaneous dissociation and ~10^6^–fold relative to RecBCD–induced dissociation of single proteins in the absence of crowding (Fig. 5B and SI Appendix, Fig. S7)(19). A representative MD simulation trajectory of RecBCD traveling on DNA bound by three EcoRI^E111Q^ molecules is shown in Fig. 5C and *SI Appendix*. Remarkably, all MD simulations (N=15/15) involving RecBCD collisions with three molecules of EcoRI^E111Q^ resulted in sequential eviction of the first and then second proteins (Fig. 5D). Together, these MD simulations provide further support for a model in which RecBCD dislodges EcoRI^E111Q^ from crowded DNA by forcing the proteins into one another, leading to an accumulation of resistance that ultimately causes sequential eviction of individual proteins from within the tandem arrays.

### Protein eviction through torque-generated DNA deformation

We next sought to determine more precisely why the proteins dissociated from the DNA when forced into one another by RecBCD. Detailed analysis of the MD simulation trajectories revealed extensive DNA deformation as EcoRI^E111Q^ molecules were pushed together (Fig. 5A to D). Deformation occurs because the EcoRI center–of–mass (COM) is ~24 Å away from the DNA axis, so the force (*F*) exerted by RecBCD on EcoRI^E111Q^ when it is pushed into the distal obstacle gives rise to torque (*F* × *d*), where *d* is the distance between the DNA and the protein COM. This torque bends the DNA, which destabilizes the interface between EcoRI^E111Q^ and the DNA causing the protein to rapidly dissociate into solution (Fig. 5B and D). We conclude that EcoRI^E111Q^ dissociates from the DNA due to torque-induced deformation of the underlying protein-nucleic acid interface.

It is possible that the extensive DNA bending observed for collisions involving tandem molecules of EcoRI was a unique outcome arising because of the specific geometry of the protein-protein interface that is formed when the two molecules of EcoRI are pushed into one another. To test this possibility, we conducted MD simulations using EcoRI embedded within spherical shells. We varied the radius and COM of the spherical shells to eliminate any protein surfaces or binding geometries that may be specific to EcoRI, allowing us to emulate a series of “generic” high affinity DNA-binding proteins of varying sizes (*SI Appendix*, Fig. S8A). In all simulations, the proteins still dissociated rapidly from the DNA (*SI Appendix*, Fig. S8B) and each dissociation event was correlated with extensive deformation of the underlying protein-DNA interfaces (*SI Appendix*, Fig. S8C). Taken together, these MD simulations suggest that similar eviction mechanisms may be operative for other high affinity DNA-binding proteins, even those with different radii, different molecular interfaces or binding geometries, and different COMs. Interestingly, 87% of all protein-DNA complex structures in the protein data bank (PDB) have a COM that is offset from the DNA axis by ≥5 Å (*SI Appendix*, Fig. S8D) (32), suggesting the possibility that torque induced DNA deformation might be a general mechanism allowing RecBCD to clear DNA of crowded obstacles.

### RNA polymerase rapidly dissociates when forced into EcoRI^E111Q^

Our results demonstrate that RecBCD sequentially removes EcoRI^E111Q^ from DNA by pushing the proteins into one another. We next sought to determine whether similar principles apply to other nucleoprotein roadblocks. RNA polymerase is perhaps the most abundant and formidable nucleoprotein roadblock that will be encountered by RecBCD in living cells. A single *Escherichia coli* cell contains ~2,000 molecules of RNAP, and under typical growth conditions ≥65% of these polymerases are bound to the bacterial chromosome (33). RNAP is also of particular interest because it is a high-affinity DNA-binding protein (K_*d*_ ~10–100 pM) and a powerful translocase capable of moving under an applied load of up to ~14–25 pN (34). RNA polymerases can survive encounters with replication forks and stall fork progression in head-on collisions (35–39). Indeed, the highly transcribed ribosomal RNA genes are a potent blockade to DNA replication (40–42). We have previously shown that RecBCD can disrupt individual *E. coli* RNAP complexes, including core RNAP, RNAP holoenzyme, stalled elongation complexes and actively transcribing polymerases in either head-to-head or head-to-tail orientations (19). RecBCD pushes isolated RNAP for long distances over naked DNA, and dissociation takes place as RecBCD forces the polymerase to step from one nonspecific binding site to the next (19).

To determine whether and how RecBCD might disrupt RNAP on crowded DNA, we first sought to establish what happens when RecBCD pushes RNAP into tandem 5x arrays of EcoRI^E111Q^ (Fig. 6A). If the sequential model applies to RNAP, then this model predicts that RNAP should rapidly dissociate from the DNA upon being forced into the EcoRI array by RecBCD. For these experiments, Qdot-tagged RNAP holoenzyme was bound the native phage promoters (19, 43), and unlabeled RecBCD was loaded onto the free ends of the DNA molecules. RecBCD translocation was then initiated by the addition of ATP. Remarkably, RNAP dissociates from the DNA almost immediately upon being pushed by RecBCD into the EcoRI^E111Q^ array for all observed collisions (N=22/22) (Fig. 6A). Control experiments confirmed that RNAP dissociation at the 5x EcoRI array position was entirely dependent upon the presence of EcoRI^E111Q^, and when EcoRI^E111Q^ was absent, many of the polymerases were instead pushed to the ends of the naked DNA molecules (Fig. 6A). We conclude that RNA polymerase rapidly dissociates from the DNA when pushed into other high affinity DNA-binding proteins by RecBCD, in good agreement with the sequential eviction model.

**Fig. 6.**
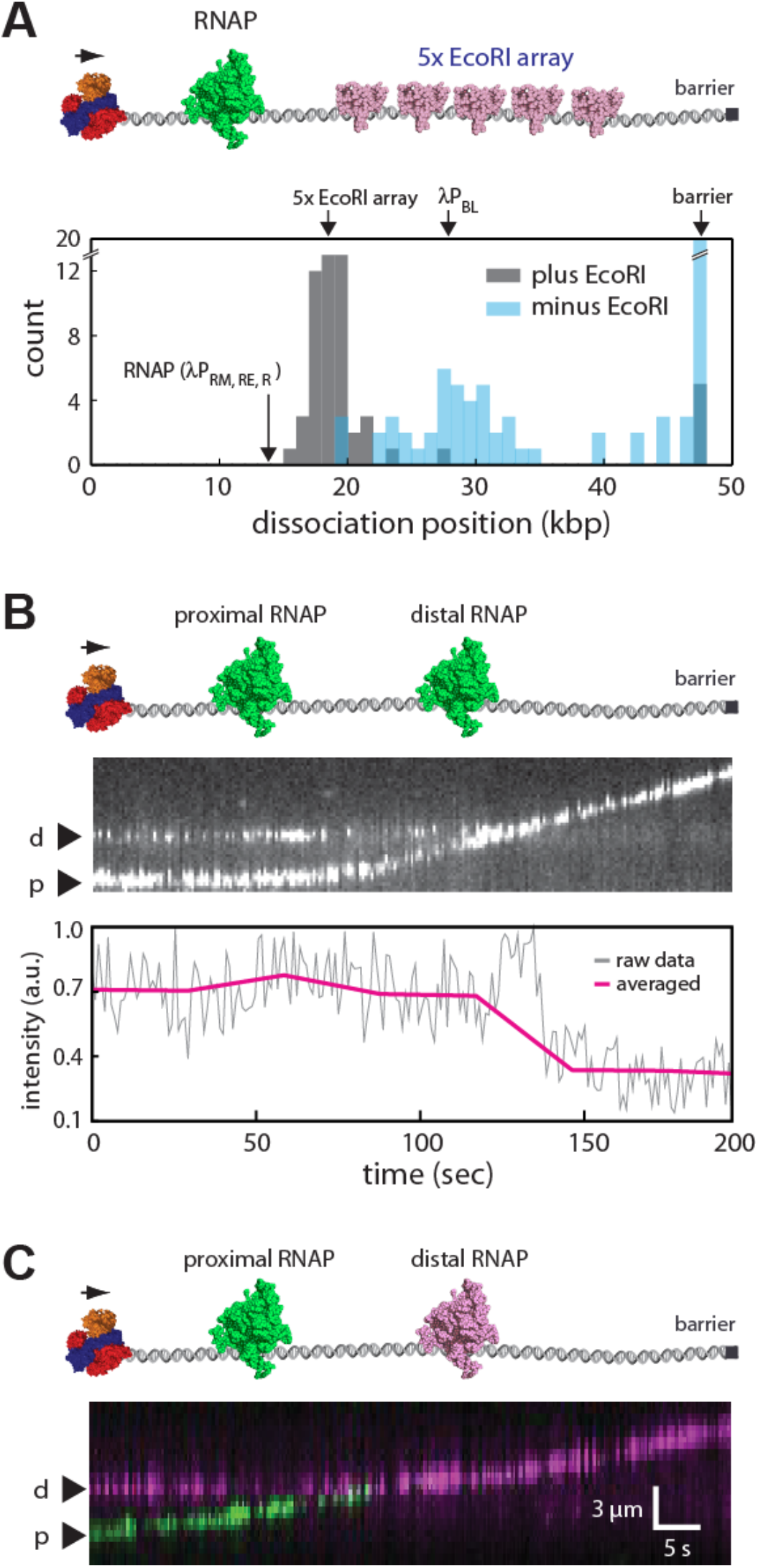
RecBCD sequentially disrupts tandem RNA polymerases. **(A)** Experimental schematic (upper panel) and resulting dissociation position distribution data (lower panel) for Qdot-tagged RNAP pushed into 5x EcoRI arrays by RecBCD. **(B)** Representative one-color kymograph showing two Qdot -tagged RNAP complexes (promoter-bound holoenzyme) being pushed into one another by unlabeled RecBCD, and the corresponding graph showing the cumulative fluorescence intensity of the Qdot-tagged proteins. **(C)** Representative two-color kymograph showing two Qdot –tagged RNAP complexes (promoter bound holoenzyme) being pushed into one another by unlabeled RecBCD; similar outcomes were observed in most (N=50/52) two–color RNAP collisions, and in the remaining cases, RecBCD pushed both proteins (N=2/52). In **(B)** and **(C)**, reactions were initiated by addition of 1 mM ATP into the RecBCD buffer (40 mM Tris–HCl [pH 7.5], 2 mM MgCl_2_, plus 0.2 mg ml^−1^ Pluronic) and gaps in the RNAP trajectories result from Qdot blinking.

### Sequential eviction of tandem RNA polymerases

Interestingly, when EcoRI^E111Q^ was absent from the reactions, a second population of RNAP dissociated at a position coinciding with the location of the λP_BL_ promoter (Fig. 6A)(43). One possible explanation for this observation is that the Qdot-tagged RNAP might be encountering unlabeled RNAP bound to the λP_BL_ promoter as it is pushed along the DNA by RecBCD, and the resulting collisions with the unlabeled proteins may have provoked rapid eviction of the Qdot-tagged protein from the DNA. Therefore, we next sought to directly examine what happens when RecBCD pushed two RNA polymerases into one another. To accomplish this, we relied upon the eight native promoters present in the λ phage genome, which allow multiple RNAP complexes to be loaded onto the same DNA molecule (43). We first performed DNA curtain experiments using unlabeled RecBCD and promoter-bound RNAP open complexes, which were labeled with a single color Qdot (Qdot 705) (Fig. 6B). Remarkably, analysis of the cumulative fluorescence intensity of the DNA-bound polymerases suggested that only one of the two polymerases remained on the DNA when RecBCD pushed them into one another (Fig. 6B). We conclude that although RecBCD readily pushes single RNAP complexes along DNA, it does not appear to push two RNAPs at the same time. Instead, as predicted by the sequential eviction model, one of the two polymerases quickly falls of the DNA when they collide with one another.

The results described above provide evidence that the sequential eviction model may apply to RecBCD encounters with RNA polymerases in crowded settings. Importantly, the sequential eviction model specifically predicts that the proximal polymerase should be preferentially evicted when pushed by RecBCD into the distal polymerase. We next sought to verify this prediction by determining which of the two polymerases dissociated from the DNA when forced into one another by RecBCD. We therefore conducted two-color DNA curtain assays in which separate aliquots of RNAP were labeled with either green (Qdot 605) or red (Qdot 705) quantum dots. The differentially labeled polymerases were then mixed together and bound to the native phage promoters, unlabeled RecBCD was bound to the free DNA ends and translocation was initiated by the injection of ATP. These experiments revealed that for ~96% of observed collisions (N=50/52) involving RecBCD and two tandem molecules of RNAP, the proximal polymerase almost immediately dissociated from the DNA upon being pushed into the distal polymerase (Fig. 6C). We conclude that RecBCD can rapidly and sequentially evict RNAP from crowded DNA, and that it does so specifically by forcing the polymerases into one another.

## Discussion

Our results demonstrate that RecBCD rapidly and sequentially removes crowded nucleoprotein complexes from crowded DNA by pushing them into one another. These findings reveal that molecular crowding itself can have a crucial and unanticipated influence on how molecular motor proteins clear nucleic acids of bound obstacles.

### Molecular crowding alters the mechanism of protein eviction by RecBCD

The mechanism by which RecBCD removes high affinity nucleoprotein complexes from DNA is dependent upon molecular crowding (Fig. 7). When RecBCD encounters isolated molecules of either of EcoRI^E111Q^ or RNA polymerase on otherwise naked DNA, it is able to push these proteins over average distances of 13,000 ± 9,100 bp and 10,460 ± 7690 bp, respectively (19). The proteins eventually dissociate as they are forced to step between successive nonspecific binding sites, and the probability (*P*) of dissociation is directly proportional to the number of steps (*n*) that EcoRI^E111Q^ or RNA polymerase are forced to take while being pushed along the DNA (Fig. 7A) (19). However, the probability of dissociating during any given step is low, and as a consequence RecBCD is able to push isolated proteins for extended distances on naked DNA (Fig. 7A). This outcome is in marked contrast to what takes place in crowded environments, where EcoRI^E111Q^ and RNA polymerase both dissociate almost immediately when pushed into more distal proteins or protein arrays (Fig. 7B). This much more rapid eviction takes place on crowded DNA specifically because RecBCD pushes the proteins into one another. Molecular dynamics simulations of collisions involving tandem molecules of EcoRI^E111Q^ suggest that when crowded proteins are forced into one another, RecBCD uses the most proximal protein as a molecular lever to generate torque, which in turn distorts the DNA and destabilizes the underlying protein-nucleic acid interface of the proximal protein (Fig. 7B, and see below). The striking differences between the outcomes of isolated collisions involving single nucleoprotein complexes, compared to the outcomes of collisions on crowded DNA, highlights the dramatic and unexpected impact that molecular crowded has on the mechanism by which RecBCD interacts with nucleoprotein obstacles that it encounters while traveling along DNA. This sequential mechanism for protein eviction may likely reflect what takes place *in vivo* where long tracts of naked DNA are unlikely to exist (44, 45).

**Fig. 7.**
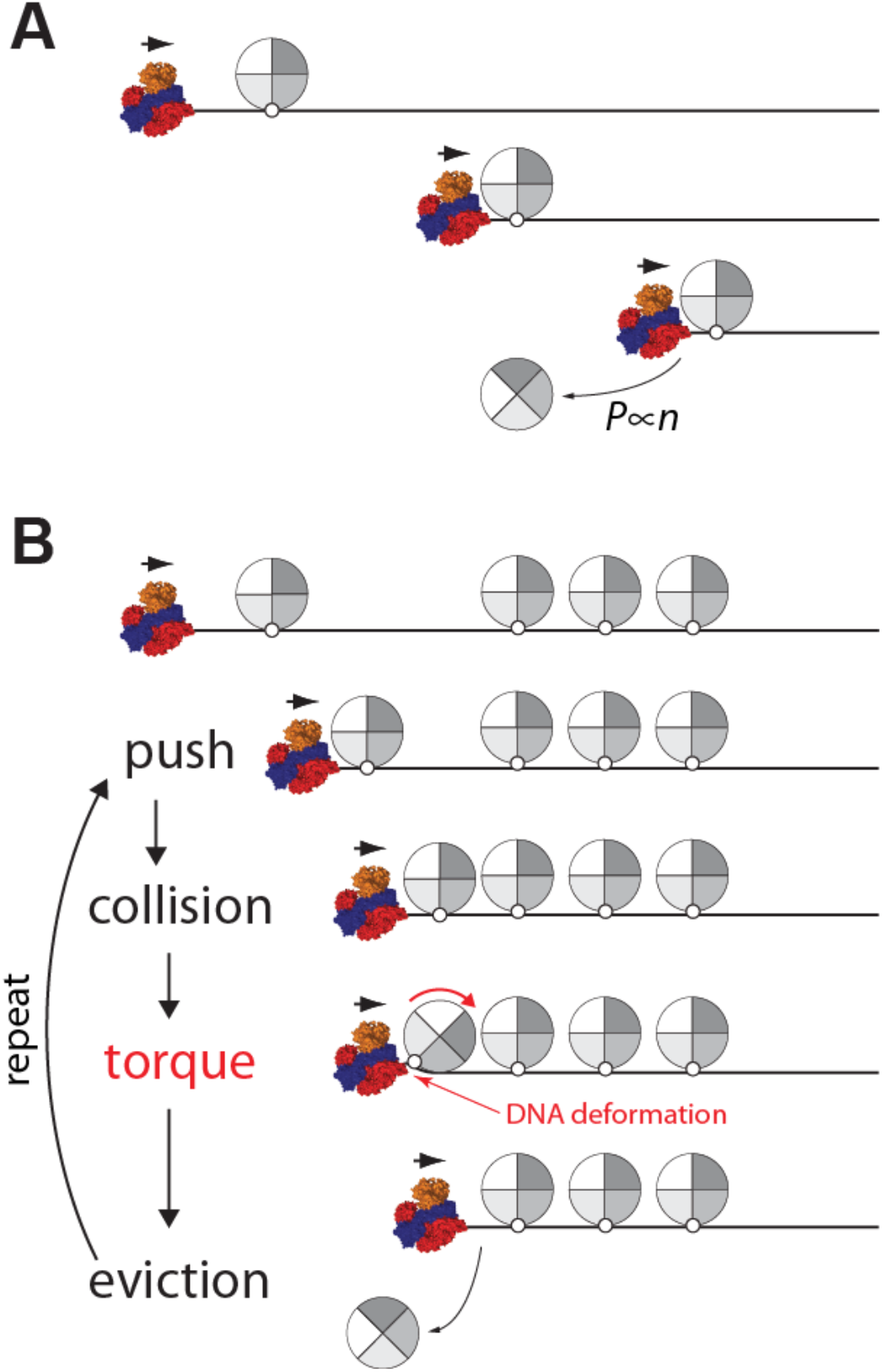
Generalized model for torque-driven protein eviction in crowded environments. **(A)** RecBCD can push isolated proteins for extended’ distances along the otherwise naked DNA, and the probability (*P*) of dissociation scales proportionally with the number of steps (*n*) that the protein is forced to take as its pushed by RecBCD (19). **(B)** In contrast, when RecBCD encounters proteins in crowded environments, it pushes each protein into its nearest adjacent neighbor. The resulting collisions generate torque on the most proximal protein to RecBCD, which distorts the DNA and disrupts the underlying protein–DNA interface resulting in rapid dissociation of the protein. Additional details of the models are presented in the main text.

### Sequential removal of RNA polymerase from DNA by RecBCD

Our model for torque-dependent sequential disruption of crowded nucleoprotein complexes has only three basic requirements: (*i*) a high-affinity DNA-bound protein whose center of mass (COM) is offset from the DNA axis; (*ii*) a motor protein capable of generating sufficient force to push the DNA-bound protein in question; and (*iii*) a downstream nucleoprotein complex to provide additional resistance. These simple requirements highlight the generality of the proposed sequential dissociation mechanism, and suggest its potential applicability to other nucleic acid motor proteins and different types of nucleoprotein obstacles. The general applicability of the sequential disruption model is also supported by the similarities between the experimental findings for EcoRI^E111Q^ and *E. coli* RNA polymerase. Molecular dynamics simulations of collisions involving RNAP are not yet feasible, so we cannot determine whether RNA polymerase dissociation also coincides with extensive deformation of nucleoprotein interface. However, the *E. coli* RNA polymerase center of mass is offset from the DNA axis by ~26.5 Å (46), suggesting that sequential eviction of RNA polymerase may also be occurring through a torque-dependent mechanism similar to that described for EcoRI^E111Q^. Future work will be necessary to more precisely determine the extent to which DNA deformation contributes to sequential disruption of RNA polymerase by RecBCD.

Interestingly, our work suggests that a small number of EcoRI^E111Q^ complexes can accumulate in front RecBCD before they start dissociating from the DNA. In contrast to EcoRI^E111Q^, it does not appear as though multiple molecules of RNA polymerase can accumulate in front of RecBCD. Instead the proximal polymerase dissociates from the DNA almost immediately upon being pushed into a distal protein. These observations indicate that RecBCD removes RNA polymerase from crowded DNA much more easily than it removes EcoRI^E111Q^. This difference may reflect the fact that RNA polymerase is a naturally occurring obstacle that will likely be encountered whenever RecBCD acts on DNA in a cellular environment. We speculate that co-evolution of RNA polymerase and RecBCD may have tuned to relative binding strengths of these two nucleoprotein complexes to ensure that RNA polymerase cannot impede the movement of RecBCD.

### Mechanisms of protein dissociation

Through-DNA allosteric communication can influence the dissociation of stationary proteins that are bound in close spatial proximity to one another (47). Previous studies have shown that protein pairs, including glucocorticoid receptor and BamHI, or lac repressor together with either EcoRV or T7 RNAP, exhibited up to 5-fold changes in dissociation rates when the corresponding partner was bound to a nearby DNA site (47). These experimental findings have been attributed to through-DNA allosteric communication based upon the long-range oscillatory changes in DNA major and minor groove widths observed in MD simulations (47–49). Similarly, we find that a static RecBCD complex positioned immediately adjacent to an EcoRI site causes approximately a 2-fold reduction in DNA cleavage by EcoRI (*SI Appendix*, Fig. S9). This reduction in cleavage is comparable in magnitude to the effects previously ascribed to the through-DNA allosteric model. However, this 2-fold effect is substantially less than the ~10^6^-fold enhancement in dissociation rates observed as RecBCD traverses an array of EcoRI^E111Q^, suggesting that through-DNA allostery may not be a predominant factor affecting protein displacement by RecBCD. In addition, our experimental data demonstrate that under crowded conditions, RecBCD causes rapid dissociation of the most proximal protein, but only when it is pushed into a more distal obstacle, indicating that crowded environments enhance protein dissociation by RecBCD relative to isolated collisions. The substantial increase in rate enhancement when RecBCD is moving through a protein array, together with the dependence of proximal protein dissociation on the presence of a more distal obstacle, suggests that the through-DNA allosteric model cannot account for RecBCD-mediated protein displacement under crowded conditions. Instead our MD simulations suggest that RecBCD can push proteins into one another, which is consistent with our experimental observations, and that the resulting collisions give rise to torque that results in rapid destabilization of the proximal protein obstacle. Although the torque-based model comes from MD simulations, the model itself is in good agreement with our experimentally observed finding that RecBCD provokes sequential protein dissociation under crowded conditions. Future work will be essential to further evaluate the torque-based model for protein eviction from crowded environments, and to determine whether other types of DNA translocases may act similarly

### Nucleoprotein obstacles and genome integrity

Nucleoprotein complexes are the primary source of replication fork stalling (50) and their presence represents a major challenge to genome integrity (1, 40, 41, 51–53). Indeed, prokaryotic and eukaryotic replisomes both require accessory helicases to clear tightly bound proteins from DNA (25, 54–58). For instance, the *E. coli* replisome requires the accessory helicases Rep and UvrD to prevent replication fork collapse upon encountering RNA polymerase and other types of high affinity nucleoprotein complexes (25, 50). Similarly, *Bacillus subtilis* requires the accessory helicase PcrA to facilitate replication fork progression through highly transcribed genes (56). The physical basis by which Rep, UvrD and PcrA assist the replisome remain unknown, however, all three proteins are 3’→5’ SF1A helicases and are closely related to RecB (13), suggesting the possibility that they may strip proteins from crowded DNA through mechanism similar to that used by RecBCD. In addition, both Rep and UvrD can push isolated ssDNA-binding proteins along single-stranded DNA (59) using a mechanism that is similar in many respects to what takes place during RecBCD collisions with isolated proteins (19). Future work will be essential for further establishing how these replication accessory helicases and other types of motor proteins disrupt tightly bound nucleoprotein complexes on crowded nucleic acids.

## Conclusion

We have presented a model describing the ability of RecBCD to sequentially clear crowded DNA of nucleoprotein complexes. The key feature of the sequential model is that RecBCD provokes rapid disruption of crowded nucleoprotein complexes by pushing these obstacles into one another. This model suggests that molecular crowding itself alters the mechanism by which RecBCD removes proteins from DNA, and the proposed mechanism is supported by a combination of experimental data, kinetic Monte Carlo simulations and molecular dynamics simulations. The sequential model, which represents the first detailed mechanistic description for the behavior of any motor protein in a highly crowded environment, strongly suggests that ATP-dependent nucleic acid motor proteins can respond differently to encounters with isolated nucleoprotein complexes compared to encounters involving multiple nucleoprotein complexes. The general principles revealed from our studies with RecBCD may also apply to behavior of other types of motor proteins as they travel along crowded nucleic acids.

## Acknowledgments

We thank members of the Greene laboratory for discussion and assistance throughout this work, and we thank Shoji Takada, Jayil Lee, Daniel Duzdevich, Fabian Erdel, Corentin Moevus, Kyle Kaniecki, Chu Jian Ma, Johannes Stigler, Justin Steinfeld, Luisina De Tullio, and Mayu S. Terakawa for comments on the manuscript. This work was supported by a Japan Society for the Promotion of Science fellowship to T.T., an NIH Grant (R35GM118026) to E.C.G., and NIH Fellowship (F32GM101819) to T.D.S. The data described in this manuscript are archived in the laboratory of E.C.G. in the Department of Biochemistry and Molecular Biophysics, Columbia University.

## Supplemental Information Appendix

### EXTENDED EXPERIMENTAL PROCEDURES

#### Kinetic Monte Carlo simulations

The kinetic Monte Carlo simulations included three kinds of molecules: the DNA, the translocase, and the roadblock protein. The DNA was a one–dimensional track 49–kilobase pairs (kbp) in length. Within the simulations, the translocase moves along DNA at a fixed rate of 1,500 bp sec^−1^ 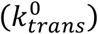. The roadblock proteins bind to and dissociate from specific binding sites with a rate constant *k^sp^* and dissociate from non–specific binding sites with a rate constant *k^ns^*. When the translocase encounters a roadblock protein, they make a complex (*tr*_1_). The *tr*_1_ complex moves along DNA with a rate constant 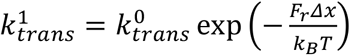, where *F_r_* is the resistance experienced by the translocase as it pushes the roadblock protein, *x* is the direction of movement, *k_B_* is the Boltzmann constant, and *T* is temperature (310 K). Likewise, when the *tr_n_* encounters the next roadblock protein, they make a larger complex (*tr*_*n*+1_). In this case, *F_r_* is dependent on the number of roadblock proteins bound to specific versus nonspecific sites such that 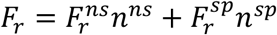, where *n^ns^* and *n^sp^* are the number of proteins bound to non–specific and specific sites, respectively. 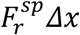 and 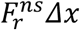 were set to 0.01 and 0.001 J mol^−1^, respectively. When the translocase encounters a roadblock protein, it can also induce the dissociation of the protein with dissociation rate constant written as 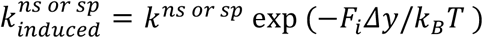, where *k^ns^* and *k^sp^* are dissociation rate constants for proteins bound to non-specific and specific binding site (set to 4.5×10^−7^ and 4.5×10^−3^ sec^−1^, respectively), *F_i_* is the force necessary for the translocase to induce protein dissociation, and *y* is the direction of dissociation.

In the initial set up, the translocase binds to one terminus of the DNA. A protein array comprised of 5–50x specific binding sites at 40–bp intervals is located 15–kbp from the DNA end. Time evolution of the system was performed using the Gillespie algorithm. In this algorithm, we first calculate escape rate constant 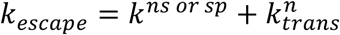. Then, we calculate the time when a particular event takes place (*τ* = −1/*k_escape_logU*; *U* is random number; 0 < *U* < 1) as well as the probability of each particular event ( *p_event_* = *k_event_*/*k_escape_*). The current time and state of the system are updated accordingly and the procedure is repeated until the translocase reaches the opposite DNA end. The three different scenarios illustrated in Fig. 1A were realized by altering the parameter values for 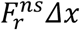, 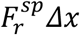, *k^ns^* and *k^sp^*. For the accumulation model, we prohibit the protein dissociation by setting *k^sp^* = 0 and *k^ns^* = 0. For the sequential model, we ensured more rapid protein dissociation upon collision with the translocase by setting *F_i_Δy* to 0.4 J mol^−1^. For the spontaneous dissociation model, we set 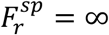 and 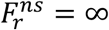 to prohibit further movement of the translocase complex upon encountering a protein until the protein spontaneously dissociates from the DNA. In this spontaneous dissociation model, the average pause duration can be analytically derived and as < *t* >≈ *k*^−1^ log(*N*_0_), where *k* is the rate constant for dissociation, and N(t) and N_0_ are the number of proteins bound to the array initially and at time *t*, respectively, in accord with the simulation result.

For the alternative variation of the accumulation model, we varied the parameter describing resistance of a roadblock protein on specific binding sites 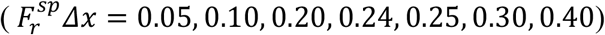 while leaving the other model parameters at fixed values 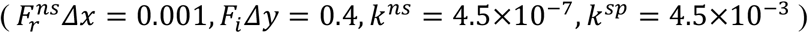. This setting allows us to change the relative rates of EcoRI^E111Q^ dissociation and sliding, resulting in the accumulation of varying numbers of EcoRI^E111Q^ in front of RecBCD prior to dissociation (*SI Appendix*, Fig. S1A). Each different value of 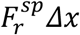, leads to the accumulation of a different number (*N*) of EcoRI^E111Q^ proteins in front of RecBCD (*SI Appendix*, Fig. S1B).

#### Proteins and DNA

Biotinylated RecBCD (RecBCD–biotin) was purified from *E.coli* JM109(DE3) cells co– transformed with plasmids for expression of RecC and RecBD with an avidity tag on the C– terminus of RecD. The cells were grown in 2YT to *OD*_600_~0.6 in the presence of 25 μg ml^−1^ Chloramphenicol and 100 μg ml^−1^ Carbenicillin. The media was then supplemented with 0.2 mM biotin and protein expression induced by addition of 0.5 mM IPTG, and cells were grown for an additional 16 hours at 16°C. Cells were then harvested by centrifugation and re–suspended in lysis buffer (50 mM Tris–HCl [pH 7.5], 0.5 mM PMSF, 10% sucrose) and lysed by sonication. The lysate was clarified by centrifugation and the supernatant was loaded onto the chitin column (New England Biolabs). The column was washed with buffer A (40 mM Tris–HCl [pH 7.5], 100 mM NaCl, 1 mM EDTA, 5% Glycerol) and RecBCD was eluted with buffer B (40 mM Tris–HCl [pH 7.5], 100 mM NaCl, 1 mM EDTA, 5% Glycerol, 50 mM DTT). Following the purification, RecBCD was dialyzed into storage buffer containing 40 mM Tris–HCl [pH 7.5], 100 mM NaCl, 1 mM EDTA, and 35% Glycerol, then frozen on liquid nitrogen and stored at –80°C.

Catalytically inactive EcoRI^E111Q^ bearing a flag epitope tag was expressed as a fusion construct linked to a self–cleaving intein and a chitin–binding domain (CBD) in *E.coli* HMS174(DE3) cells. The cells were grown in 2YT to *OD*_600_~0.6 in the presence of 100 μg ml^−1^ Carbenicillin. Protein expression was induced by the addition of 0.5 mM IPTG, and cells were grown for 4 hours at 37°C. Cells were then harvested and re–suspended in lysis buffer (50 mM Tris–HCl [pH 7.5], 0.5 mM PMSF, 10% sucrose) and lysed by freezing and sonication. The lysate was clarified by centrifugation and the supernatant was loaded onto a chitin column. The column was washed with buffer A (20 mM Tris–HCl [pH 8.5], 500 mM NaCl) and the cleaved protein was eluted with buffer B (20 mM Tris–HCl [pH 8.5], 500 mM NaCl, 50 mM DTT). The protein was dialyzed into storage buffer containing 40 mM Tris–HCl [pH 7.5], 300 mM NaCl, 10 mM 2–Mercaptoethanol, 0.1 mM EDTA, 50% glycerol, 0.15% Triton X–100, frozen on liquid nitrogen and stored at –80°C. Wild–type EcoRI was purchased from New England Biolabs.

*E. coli* RNA polymerase containing an N–terminal 6–His tag and a C–terminal HA-tagged α–subunit was purified from *E.coli* BL21(DE3) as previously described (1). Cells were grown in LB to *OD*_600_~0.6 in the presence of 100 μg ml^−1^ Carbenicillin, expression was induced by addition of 0.5 mM IPTG, and cells were grown for an additional 4 hours at 37°C. Cells were harvested and re–suspended in lysis buffer (50 mM Tris–HCl [pH 7.5], 10 mM EDTA, 5% glycerol, 1 mM DTT, 300 mM NaCl, 300 μg ml^−1^ Lysozyme) and were then lysed by freezing and sonication. The lysate was clarified by centrifugation and fractionated with the addition of 0.350 g ml^−1^ ammonium sulfate. The precipitated protein was recovered by centrifugation, re– suspended in buffer A (10 mM Tris–HCl [pH 7.5], 5% glycerol, 1 mM EDTA, 0.5 mM DTT), and loaded onto a High prep Heparin FF 16/10 column (GE Healthcare). The column was washed with buffer B (10 mM Tris–HCl [pH 7.5], 5% glycerol, 1 mM EDTA, 0.5 mM DTT, 300 mM NaCl) and then eluted with buffer C (10 mM Tris–HCl [pH 7.5], 5% glycerol, 1 mM EDTA, 0.5 mM DTT, 600 mM NaCl). The eluted protein was cleared by centrifugation and fractionated with the addition of 0.350 g ml^−1^ ammonium sulfate. The precipitated protein was recovered by centrifugation, re–suspended in buffer D (10 mM Tris–HCl [pH 7.5], 5% glycerol, 1 mM EDTA, 0.5 mM DTT, 1 M NaCl), and loaded onto a HisTrap FF column (GE Healthcare). The column was washed with buffer D and the protein was eluted with buffer E (10 mM Tris–HCl [pH 7.5], 5% glycerol, 1 mM EDTA, 0.5 mM DTT, 1 M NaCl, 500 mM imidazol). The eluted protein was diluted in the buffer A and loaded onto a MonoQ 5/50 GL column (GE Healthcare). The column was washed with the buffer F (10 mM Tris–HCl [pH 7.5], 5% glycerol, 1 mM EDTA, 0.5 mM DTT, 300 mM NaCl) and the protein was eluted with a linear gradient to buffer G (10 mM Tris– HCl [pH 7.5], 5% glycerol, 1 mM EDTA, 0.5 mM DTT, 500 mM NaCl). Purified RNAP was dialyzed into storage buffer (50 mM Tris–HCl [pH 7.5], 50% glycerol, 1 mM EDTA, 1 mM DTT, 250 mM NaCl) and stored at –80°C.

A synthetic DNA fragment bearing five EcoRI binding sites (5’–GAATTC–3’) at 40–bp intervals was cloned into the SpeI and SalI sites of pUC19, and this plasmid (pUC19–5xEcoRI) was used a starting point for constructing the larger arrays using a strategy similar to that originally used for generating long arrays of lac operator sites (2). The 5x array fragment has a single XbaI site near one end, and the 5xEcoRI fragment was excised by digestion with XbaI and SpeI and purified by agarose gel electrophoresis. A second aliquot of pUC19–5xEcoRI plasmid was linearized with XbaI and SalI and the pUC19 backbone containing 5xEcoRI binding site was also purified. Then, these two purified DNA fragments were ligated together to generate pUC19– 10xEcoRI. Repetition of this procedure was used to generate pUC19–30xEcoRI and pUC19– 50xEcoRI. The EcoRI arrays were then excised by digestion with SpeI and SalI and then cloned into the NheI and XhoI within the λ genome. The resulting ligation products were packaged into phage particles using Gigapack III packaging extracts (Agilent Technologies; Cat. No. 200201), and the resulting phage DNA was purified as previously described (1). All the restriction enzymes and ligases were purchased from New England Biolabs.

#### Single molecule imaging

Single molecule experiments were conducted using a custom–built total internal reflection microscope (TIRFM) and DNA curtains, as previously described (3, 4). In brief, a biotinylated λ DNA molecules were anchored to a supported lipid bilayer on the surface of a microfluidic sample chamber through a biotin–streptavidin linkage, as previously described (3, 4). The DNA molecules were then pushed by buffer flow into nanofabricated chromium barriers, which disrupt the bilayer and allow the anchored DNA molecules to align with one another on the sample chamber surface.

For RecBCD experiments, the sample chambers were pre–equilibrated with RecBCD buffer (40 mM Tris–HCl [pH 7.5], 2 mM MgCl_2_, 0.2 mg ml^−1^ Pluronic). The sample chamber was then blocked by the addition of RecBCD buffer supplemented with 10 μM free biotin. Biotinylated RecBCD was labeled with streptavidin–coated 705 quantum dots (Qdots; Thermo Fisher Scientific; Cat. No. Q10163MP). Qdot −tagged RecBCD (10 nM) in 100 μl of RecBCD buffer was then injected into the sample chamber and incubated 20 minutes to allow binding. Free RecBCD was then flushed from the sample chamber and translocation was initiated by the addition of RecBCD buffer supplemented with 1 mM ATP while collecting images at 5.0 Hz. All RecBCD experiments were conducted at 37°C. We have previously reported a single pre-Chi RecBCD velocity of 1,484±167 bp/sec based on N=100 molecules of RecBCD (3). Here we find two populations with velocities of 745±37 bp/sec (corresponding to the expected velocity of RecB) and 1368±18 bp/sec (corresponding to the expected velocity of RecD) based upon N=269 molecules of RecBCD. We believe that the second slower population is now observable within the data simply because of the larger number of experimental observations, and we suspect that this small population reflects a subset of complexes in which RecD is inactivated. Therefore, we only provide a collision analysis of RecBCD complexes that exhibited translocation velocities of >1000 bp/sec. In future work we hope to more fully analyze the slower population, as well as a detailed analysis of pre- and post-chi behaviors.

#### Unlabeled EcoRI^E111Q^ experiments

For assays using Qdot–RecBCD and unlabeled EcoRI^E111Q^ (Fig. 2), the λ DNA substrates (150 pM) bearing different size EcoRI arrays were pre–incubated with EcoRI^E111Q^ before injecting the DNA into the sample chambers. The amount of EcoRI^E111Q^ used in the assays was sufficient to saturate each array, and the amount of protein utilized in each experiment scaled with the size of the array as follows: 5x array, 5 nM EcoRI^E111Q^; 10x array, 10 nM EcoRI^E111Q^; 30x array, 30 nM EcoRI^E111Q^, and 50x array, 50 nM EcoRI^E111Q^. All pre–incubations were conducted in 100 μl of EcoRI buffer (40 mM Tris–HCl [pH 7.5], 150 mM NaCl, 10 mM MgCl_2_, and 0.2 mg ml^−1^ Pluronic). The binding reactions were then diluted into 1 ml of RecBCD buffer, injected into the flowcell, and incubated for an additional 30 minutes. Free DNA and excess EcoRI^E111Q^ were then flushed from the sample chambers using RecBCD buffer. Qdot–tagged RecBCD was then injected into the sample chambers and translocation was initiated by the addition of 1 mM ATP, as described above.

#### Single–color EcoRI^E111Q^ experiments

For DNA curtain assays using unlabeled RecBCD and Qdot–tagged EcoRI^E111Q^ (Fig. 4 and 5), the unlabeled EcoRI^E111Q^ was pre–incubated with the λ DNA and then injected into the flowcell as described above. The DNA–bound EcoRI^E111Q^ molecules were then labeled with anti–FLAG antibody conjugated Qdots by injecting the Qdots (10 nM) into the flowcell, followed by a 5– minute incubation before flushing the unbound Qdots from the sample chamber. The antibody conjugated Qdots were prepared using the SiteClick™ Qdot^®^ Antibody Labeling Kit (Thermo Fisher Scientific; Cat. No. S10469 for Qdot 605 and Cat. No. S10454 for Qdot 705).

#### Two–color EcoRI^E111Q^ experiments

λ DNA (150 pM) with a 5x EcoRI binding site array was pre–incubated with 5 nM EcoRI^E111Q^ in 100 μl of EcoRI buffer (40 mM Tris–HCl [pH 7.5], 150 mM NaCl, 10 mM MgCl_2_, and 0.2 mg ml^−1^ Pluronic). The reaction was then diluted to a total volume of 1 ml in RecBCD buffer, injected into a flowcell, and incubated for an additional 30 minutes. Free DNA and EcoRI^E111Q^ were then flushed out of the sample chamber. The DNA–bound EcoRI^E111Q^ was then labeled by injecting anti–FLAG antibody conjugated Qdots (5 nM) with different emission maxima (605 nm or 705 nm, colored green and magenta in all kymographs) followed by a 5 minute incubation. RecBCD was then loaded onto the DNA, translocation was initiated by the addition of 1 mM ATP, and data collected as described above. Note that the binding distributions of “magenta” and “green” EcoRI^E111Q^ were random, and reaction trajectories in which the two colors were appropriately segregated between the 5x arrays and 1x native binding sites were selected by visual inspection (Fig. 5).

#### Experiments with RNA polymerase

Wild-type λ DNA (150 pM) was diluted to a total volume of 1 ml in RNAP buffer (40 mM Tris– HCl [pH 7.5], 100 mM KCl, 2 mM MgCl_2_, 0.2 mg ml^−1^ Pluronic), injected into the flowcell, incubated for 30 minutes, and then free DNA was flushed away. HA–tagged RNAP (500 pM) was then injected into the sample chamber in RNAP buffer and incubated for 10 minutes. The DNA–bound RNAP was labeled by injection of anti–HA conjugated Qdots (10 nM). For single–color experiments RNAP was labeled with only one color Qdot (Qdot 705). For two–color experiments the labeling reaction included an equimolar mixture of two different colored Qdots (Qdot 705 and Qdot 605) (Fig. 6). Heparin (0.2 mg ml^−1^) was included in the labeling buffer to ensure removal of any nonspecifically bound RNAP and reactions were incubated for 5 minutes. RecBCD was then loaded onto the DNA, translocation was initiated by the addition of 1 mM ATP, and data was collected as described above.

#### Coarse–grained molecular dynamics simulations

We utilized the one–bead–per–one–amino–acid model for proteins (AICG2 model (5)), and the three–beads–per–one–nucleotide model for DNA (3SPN.2 model (6)); the AICG2 model stabilizes the native protein conformation and the 3SPN.2 model stabilizes the B–type DNA conformation (5, 6). The parameters of used within these models were separately calibrated in independent studies to reproduce the physicochemical properties of these biomolecules (5, 6), and there were no other additional free parameters required except for the binding affinity of EcoRI for DNA, which was calibrated to reproduce the experimental dissociation constants (see below).

The interaction between EcoRI^E111Q^ and the DNA was modeled by the equation:

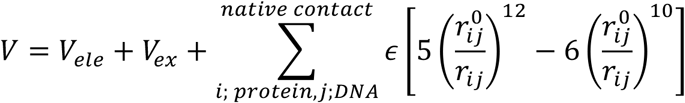

In this equation, *V_ele_* and *V_ex_* are potential energy functions for electrostatic interaction and excluded volume effect, respectively. The summation is taken for protein particles *i* and DNA particles *j* that make a contact in the crystal structure (PDB ID: 1ER1)(7). We consider two particles to make a contact when one of the heavy atoms represented by the *i*–th particle is within 6.5 Å from those represented by the *j* −th particle. The parameter 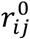 is the distance between the two particles within the crystal structure. We assumed that the EcoRI–DNA complex has indiscernible conformations on the specific and the non–specific binding site that give rise to different interaction strengths. *ϵ* is a global scaling parameter, which describes the binding energy of EcoRI^E111Q^ in association with its specific and nonspecific binding sites. To select appropriate values for *ϵ*, we first performed an umbrella sampling simulation of the EcoRI^E111Q^–DNA complex using the crystal structure (PDB ID: 1ER1) as an initial conformation and the distance between EcoRI and the DNA as a reaction coordinate (8). From this simulation we obtained a free energy curve (*SI Appendix*, Fig. S4), which was then used to calculate the free energy difference (Δ¾ between the bound and unbound states. The values for ΔG were varied by changing *ε*, allowing us to select values of *ϵ* that recapitulated the known dissociation rate constants for EcoRI^E111Q^ bound to either specific or nonspecific sites. Based on this analysis we selected *ϵ* = 0.150 for specific and *ϵ* = 0.125 for non–specific binding sites.

The initial DNA structures bound by either two (Fig. 3A and *SI Appendix*) or three (Fig. 3C and *SI Appendix*) molecules of EcoRI were built by linking EcoRI–DNA structures (PDB ID: 1ER1) to 40–bp DNA fragments, yielding final DNA lengths of 100–bp or 150–bp for the two and three EcoRI arrays, respectively. RecBCD wraps around DNA (9), and we make the simplifying assumption that the leading edge of RecBCD has pseudo-rotational symmetry about the DNA axis. Therefore, RecBCD was modeled as a simple toroidal wall with a radius of 50 Å and a hole radius of 15 Å. Treating RecBCD as a torus simplified the computational work, but does allow analysis of potential effects from protein-specific geometries; future work with more detailed models will be essential to determine whether protein-specific geometries influences the collision outcomes. One terminus of the duplex DNA was threaded through the torus in the initial structure and the DNA was then pulled through the torus to mimic RecBCD translocation along the DNA while exerting 100 pN of force. The helicase activity of RecBCD was emulated by pulling the termini of the two single strands of DNA at angle of +75° and –75° relative to the helical axis of the DNA duplex. We selected 100 pN because this value was within an order of magnitude of the stall force for several well characterized motor proteins (10). All particle coordinates were continually updated throughout the simulation according to the Langevin equation. For each setup we conducted 20 simulations by Langevin dynamics for 5–μs with friction coefficient of 0.02 ps^−1^, and temperature of 300 K. All simulations were performed using CafeMol (11). Based upon comparisons of the protein diffusion coefficients within the coarse–grained simulations with the values calculated from the Stokes–Einstein equation, we conclude that each MD simulation step corresponds to ~1 ps, as previously described (12).

For MD simulations involving RecBCD collisions with two molecules of EcoRI^E111Q^, 15 out of 15 successful simulation runs resulted in rapid dissociation of the proximal EcoRI^E111Q^ protein (protein “A” in Fig. 3A) upon colliding with the distal protein (protein “B” in Fig. 3A). For MD simulations involving RecBCD collisions with three molecules of EcoRI^E111Q^ resulted in sequential eviction of the first (protein “A” in Fig. 3C) and then second proteins (protein “B” in Fig. 3C) in 15 of 15 simulations. In 11 of these 15 three protein simulations, proteins A and B dissociated sequentially after being pushed into protein C. In the remaining 4 MD simulations, protein A dissociated upon being pushed into protein B, and then protein B dissociated upon encountering protein C.

#### Collision–induced rate enhancement

To calculate the dissociation rate constant of EcoRI^E111Q^ from non–specific DNA, we performed simulations in which a force (0.15, 0.16, 0.17, or 0.20 pN) was applied on the DNA perpendicular (*F*_⊥_) to the helical axis and away from the bound protein (*SI Appendix*, Fig. S7). For these simulations we used the EcoRI crystal structure (PDB ID: 1ER1) as an initial structure. Within the simulations each EcoRI atom was fixed in space using harmonic potentials with a spring constant of 0.05 kcal mol^−1^ Å^−2^. By extrapolating the rate constants derived from these simulations, we calculated the dissociation rate constant of EcoRI from non–specific DNA, which we then compared to the experimental dissociation rates measured EcoRI^E111Q^ either with or without RecBCD collisions (*SI Appendix*, Fig. S7B).

#### DNA bending analysis

To assess the structural distortion of the DNA as a consequence of the protein-protein collisions we defined a bending score as (*s*) = (*x*_*i*+10_ – *x_i_*)×(*x_i_* – *x*_*i*−10_), where *x_i_* is the location of the center of mass (COM) of the *i^th^* base pair. The DNA bending scores were then calculated for each simulation trajectory and used to describe the extent of structural distortion that coincided with eviction of EcoRI^E111Q^ from the DNA as a consequence of the force applied by RecBCD.

#### Shell encased collisions

To determine whether MD simulation reflected specific behavior of only EcoRI we encased the simulated EcoRI proteins within spherical shells of varying sizes. These encased molecules were intended to emulate “generic” DNA–binding proteins by eliminating any results that might require a specific protein–protein interface during the collision events. The COM of the shell was either centered over the entire EcoRI dimer COM, or the COM defined as proximal to the DNA (proximal site; located 4.64 Å away from the DNA axis), or the COM defined as distal to the DNA (distal site; located 37.57 Å away from the DNA axis) (*SI Appendix*, Fig. S8). The proximal and distal COMs were computationally treated as the averaged positions of 141^st^ and 226^th^ amino acid positions within the EcoRI dimer, respectively. The radius of the shell was 40 Å unless otherwise stated.

#### Through-DNA allosteric inhibition by RecBCD

Previous studies have demonstrated that through-DNA allosteric communication can reduce the affinity of site-specific DNA-binding proteins by up to a factor of 5 (13). Therefore, we devised a bulk biochemical measurement to determine whether (static) RecBCD might inhibit the binding of EcoRI to a nearby site, which would be consistent with the through-DNA allosteric model (*SI Appendix*, Fig. S9). For this, we designed hairpin oligonucleotides labeled with Alexa888 for detection. The oligonucleotides contained a free blunt end for binding RecBCD and a single EcoRI cleavage located either immediately adjacent to RecBCD (proximal) or separated by a 10-bp spacer from the RecBCD binding site (distal). The oligonucleotides were incubated at 95°C for 10 min, and then cooled at room temperature for 1 hour. The annealed oligonucleotides were then pre-incubated for 10 minutes with RecBCD in 18 μL of buffer containing 40 mM Tris-HCl (pH 7.5) and 2 mM MgCl_2_. Cleavage reactions were initiated by the addition of 2.0 μL of 10 μM wild type EcoRI. Aliquots (2 μL) were then transferred new tubes containing termination buffer (90% form amide, 0.5% EDTA, and 0.1% orange G) at the indicated time intervals. Samples were resolved on 10% denaturing urea polyacrylamide gels at 200V for 50 minutes. The Alexa488 signal was detected with a Typhoon FLA 7000 (GE Healthcare) and images analyzed using ImageJ (https://imagej.nih.gov/ij/). Data were analyzed by plotting the normalized signal intensity of the 30 minutes minute time points in the presence and absence of RecBCD. The results of these experiments show that DNA cleavage by EcoRI slows in the presence of RecBCD and the effect is greater for the proximal versus distal EcoRI substrate, and the relative magnitude of these effects are consistent with previous studies of through-DNA allosteric inhibition (13).

### SUPPLEMENTAL FIGURE LEGENDS

**Fig. S1.**
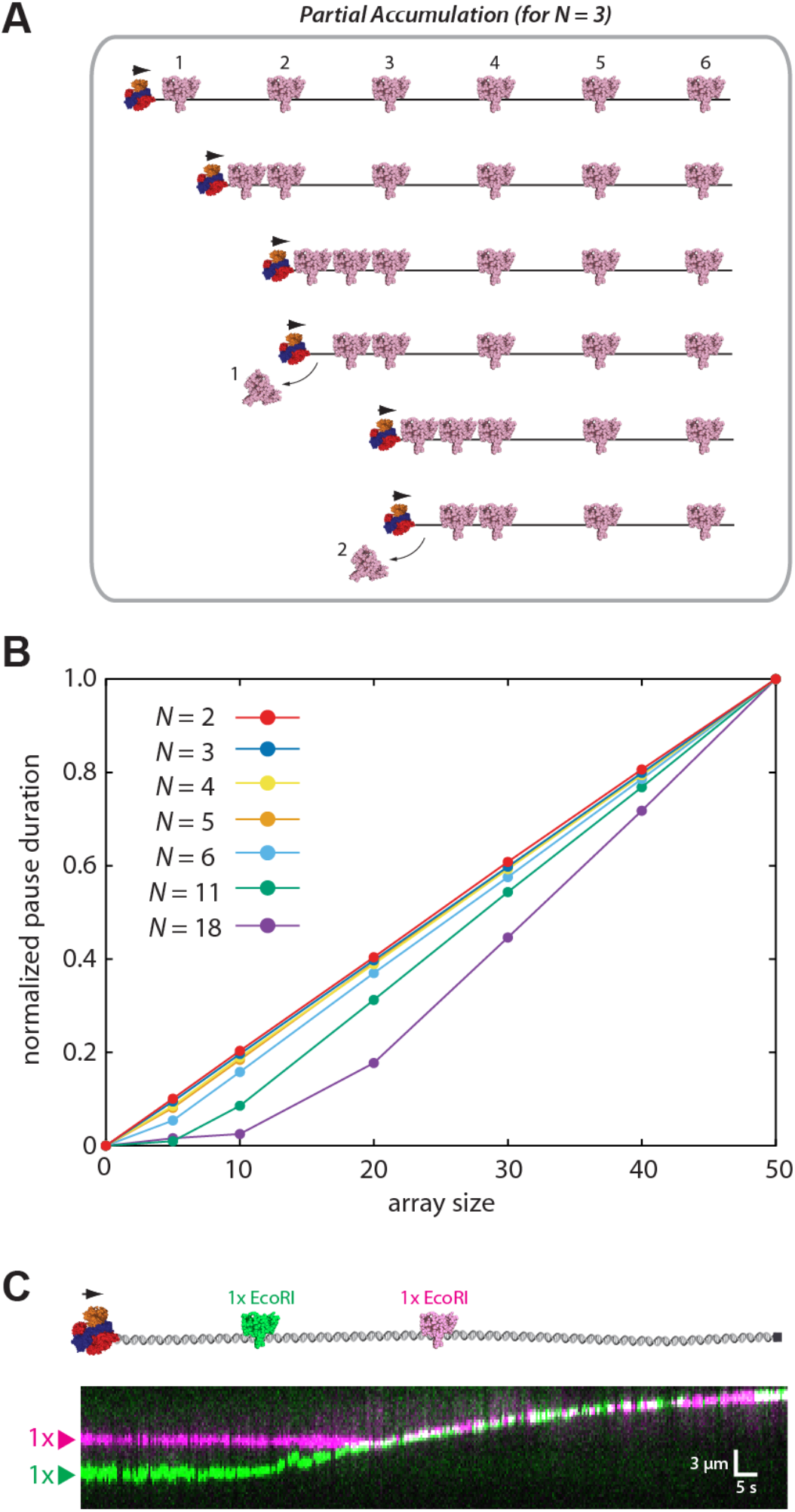
Kinetic Monte Carlo simulations of the alternative sequential dissociation model. **(A)** Schematic illustration of a hybrid model invoking components of both accumulation and sequential eviction. In this example three proteins accumulate (*N* = 3) prior to dissociation. Additional details of this model are presented in the Extended Experimental Procedures. **(B)** Pause duration distributions (〈Δt〉) for simulations involving differential numbers of accumulated proteins (*N*) for each different array size, as indicated. **(C)** Experimental schematic and kymograph demonstrating that RecBCD (unlabeled) can push two molecules of differentially labeled EcoRI^E111Q^.

**Fig. S2.**
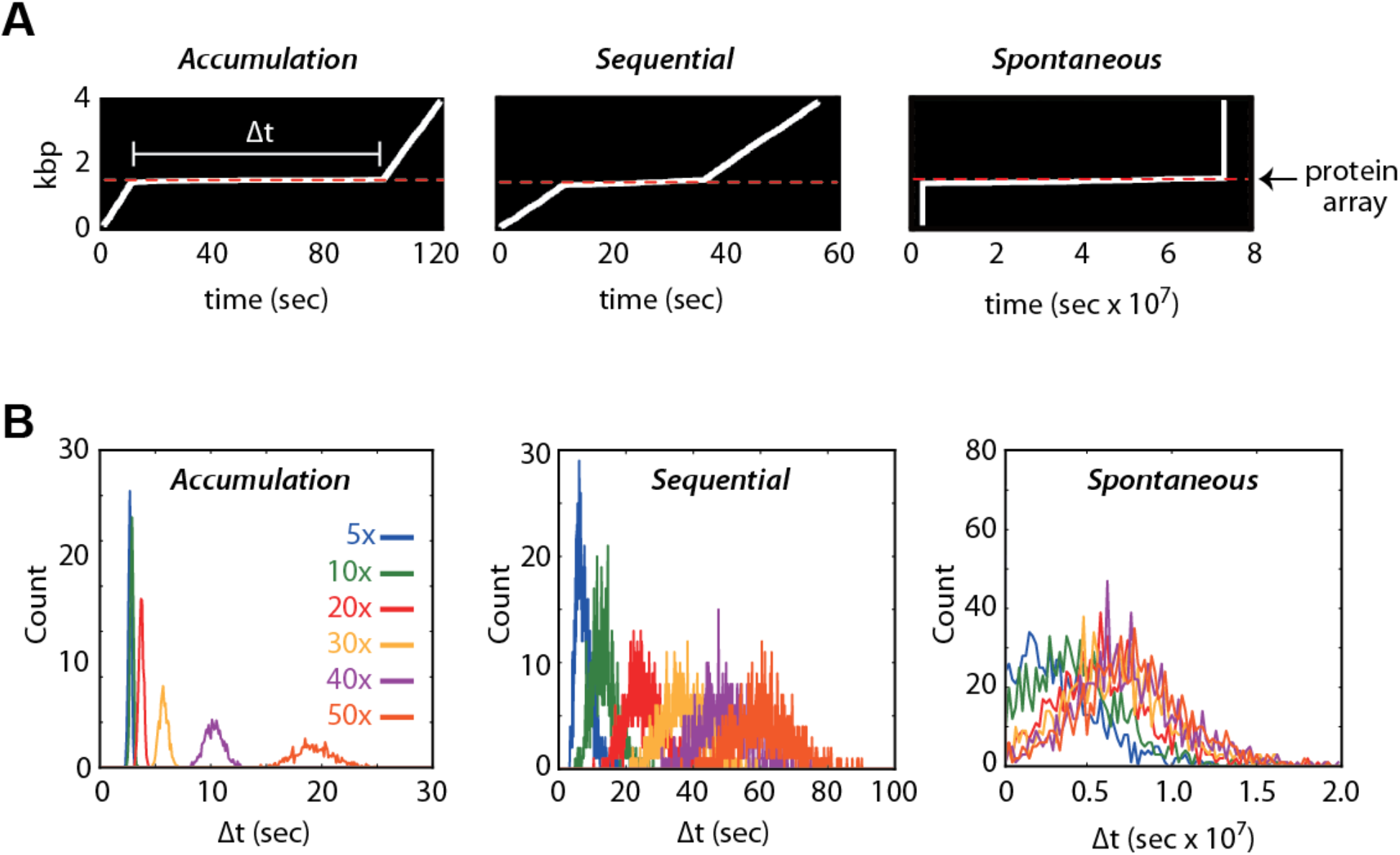
Kinetic Monte Carlo simulations to predict translocase movement through protein arrays. **(A)** Representative kymographs derived from the kinetic Monte Carlo simulations of the accumulation, sequential, and spontaneous models, respectively. The white lines represent the movement of the translocase along a DNA molecule and then pausing upon encountering the protein array. **(B)** Distributions of pause durations (〈Δt〉). Colored lines represent the distribution derived from simulations using the 5–50x protein arrays, as indicated. Each distribution was calculated from a total of 1000 simulation trajectories. **(C)** Predicted post–array RecBCD velocities for each different model.

**Fig. S3.**
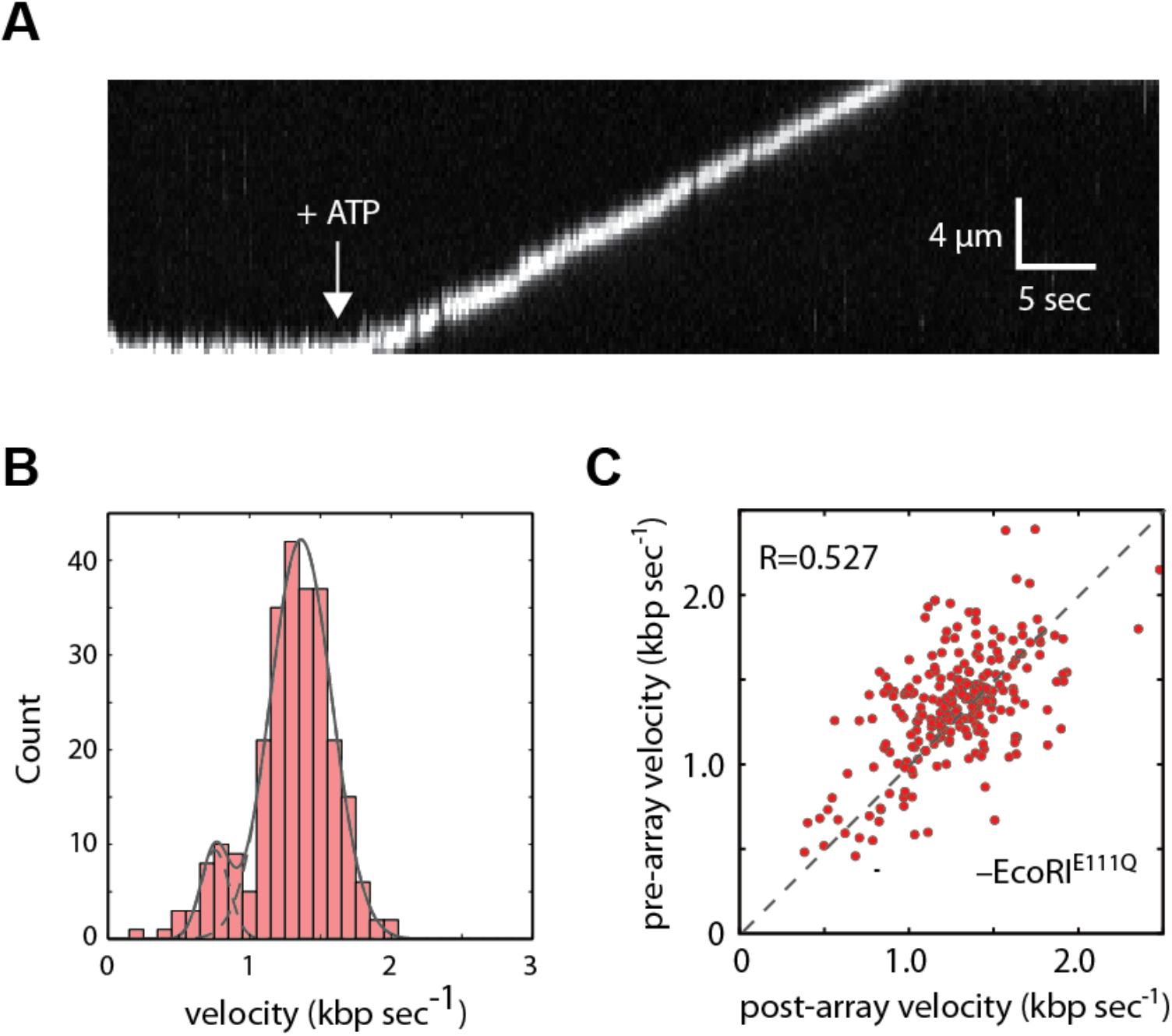
Translocation properties of Qdot–tagged RecBCD in the absence of EcoRI^E111Q^. **(A)** Example of a kymograph showing the movement of Qdot–tagged RecBCD along an unlabeled DNA molecule in the absence of EcoRI^E111Q^. RecBCD was pre–bound to the DNA and translocation initiated by the injection of ATP (white arrowhead). **(B)** Velocity distribution for Qdot–tagged RecBCD (N=269). **(C)** Scatter plot of RecBCD velocities before and after encountering the EcoRI binding site array for control experiments conducted in the absence of any EcoRI^E111Q^ (N=269).

**Fig. S4.**
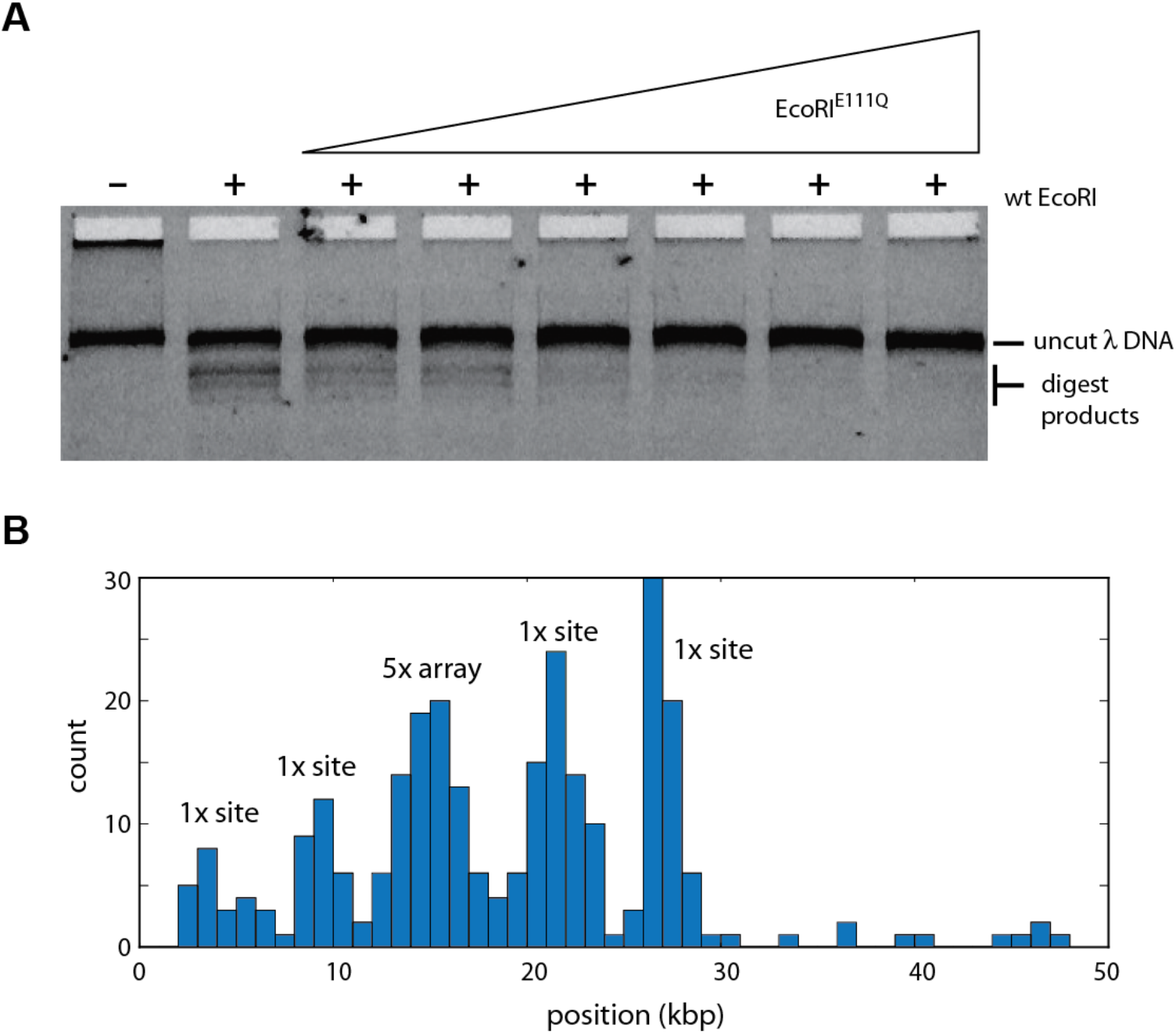
The saturation of the λ–DNA substrate with EcoRI^E111Q^. **(A)** Example of the bulk biochemical assay used to establish conditions for saturating the EcoRI binding site arrays with EcoRI^E111Q^. In this example, the λ–DNA substrate bearing a 5x binding site array was pre– incubated with varying concentrations of EcoRI^E111Q^ for 30 min at room temperature. The reactions were then challenged by the addition of wild–type EcoRI for 15 min at 37°C. DNA products were then resolved on a 1% agarose gel. **(B)** Position distribution histogram from a DNA curtain assay with Qdot–tagged EcoRI^E111Q^ bound to a λ–DNA harboring a 5x array. The location of the array is indicated, as are the four peaks corresponding to the native EcoRI binding sites.

**Fig. S5.**
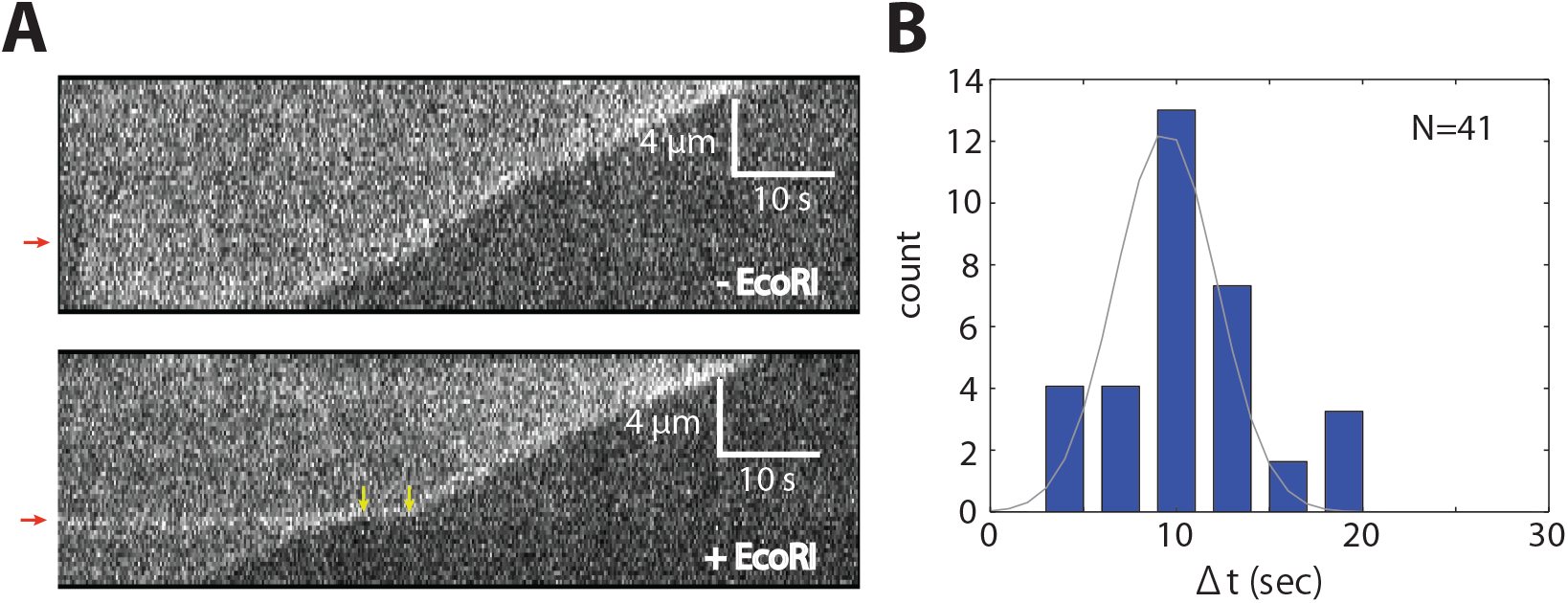
Collision of unlabeled RecBCD with an unlabeled 5x EcoRI array. **(A)** Representative kymographs from experiments where unlabeled RecBCD translocates on YoYo-1 stained DNA containing the 5x EcoRI binding site array (position indicated by red arrows) in the absence (top) and presence (bottom) of unlabeled EcoRI^E111Q^. The yellow arrows in the lower kymograph indicate the beginning and ending of the RecBCD pause when EcoRI^E111Q^ is bound to the DNA. Note that the protein-bound EcoRI arrays routinely fluoresce more brightly with YoYo-1 compared to the naked flanking DNA. **(B)** Pause time distribution for unlabeled in the presence of unlabeled EcoRI^E111Q^. The data set is fitted by a Gaussian distribution to derive the average pause duration (9.5 ± 0.4 sec).

**Fig. S6.**
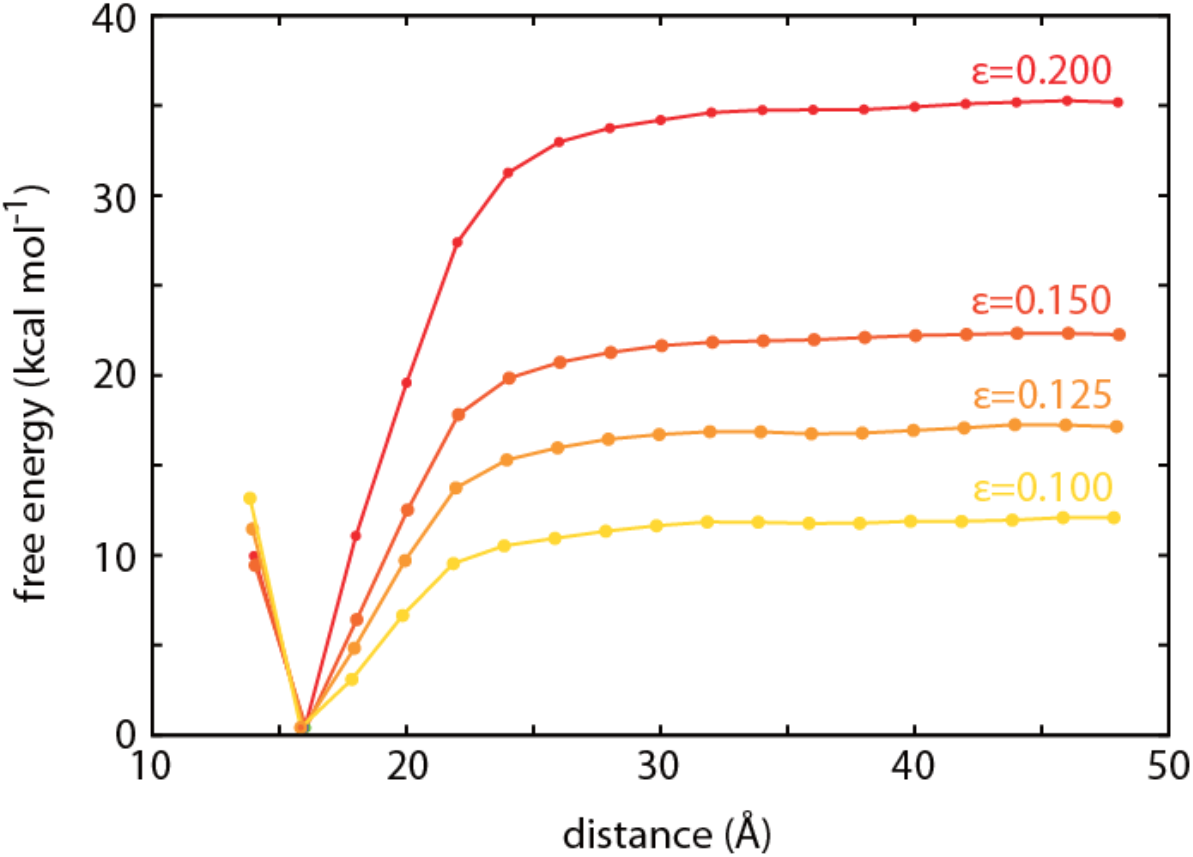
Simulation results recapitulating EcoRI^E111Q^ specific and nonspecific DNA binding characteristics. Free energy curves along the distance between EcoRI and the DNA derived from the umbrella sampling simulations (8). The different lines represent the free energy surfaces derived from molecular dynamics simulations performed for different values of the global scaling parameters (*ϵ*), as indicated.

**Fig. S7.**
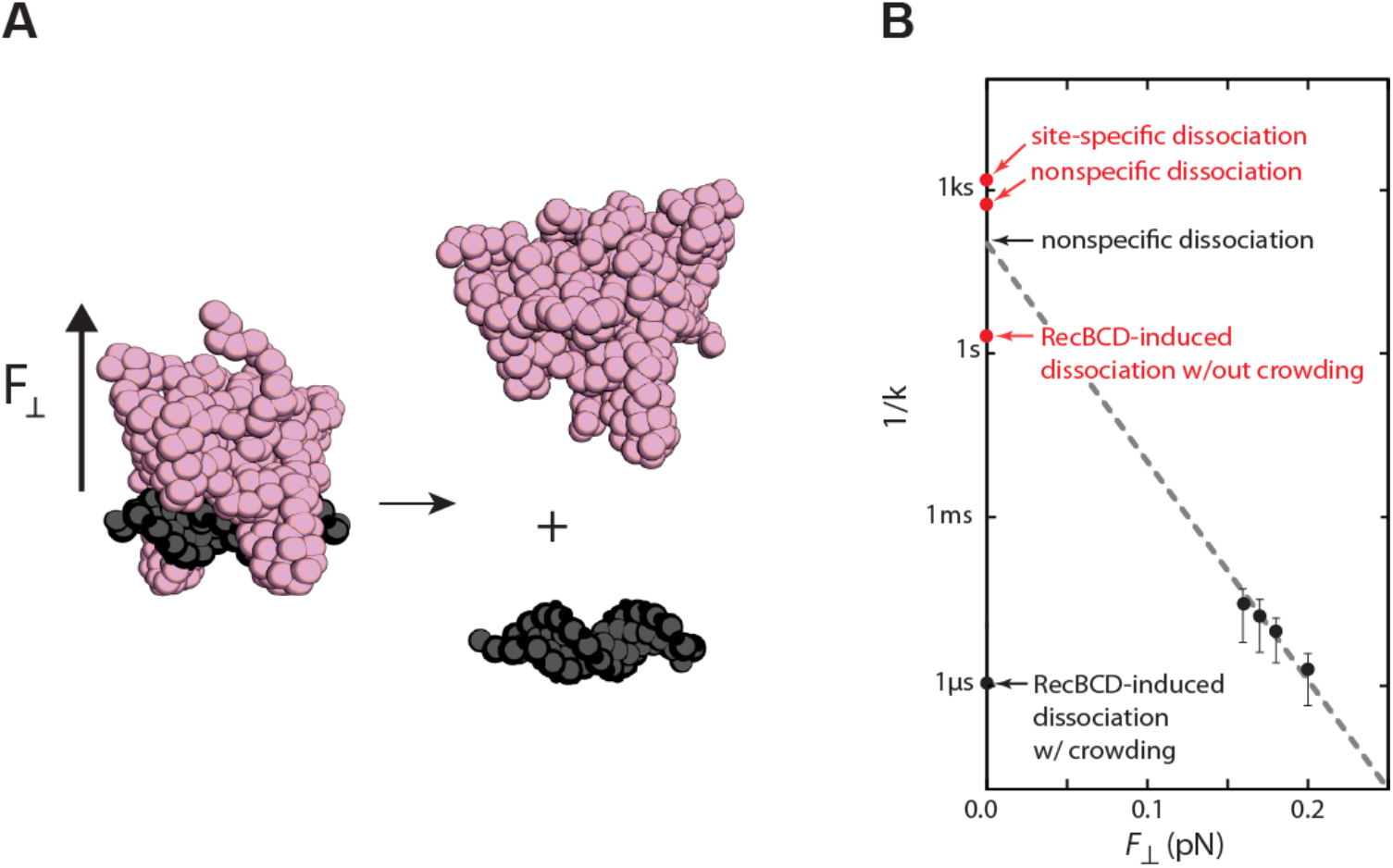
EcoRI^E111Q^ dissociation rates. **(A)** Illustration showing the application of a perpendicular force (*F*_⊥_) relative to the DNA axis, which was used to calculate EcoRI^E111Q^ dissociation rates from nonspecific DNA. **(B)** EcoRI^E111Q^ dissociation rate constants derived from the collision simulation and the simulation in which the dissociation is accelerated by application of a perpendicular force *F*_⊥_, as indicated (black). We also plotted previously reported experimental values for RecBCD collisions with single molecules of EcoRI^E111Q^ and EcoRI^E111Q^ dissociation in the absence of RecBCD (3), as indicated (red). Extrapolation of the values from the accelerated simulation lead to a nonspecific dissociation rate constant of 0.007±0.004 sec^−1^, which was in good agreement with experimental observations (0.002±0.001 sec^−1^) (3).

**Fig. S8.**
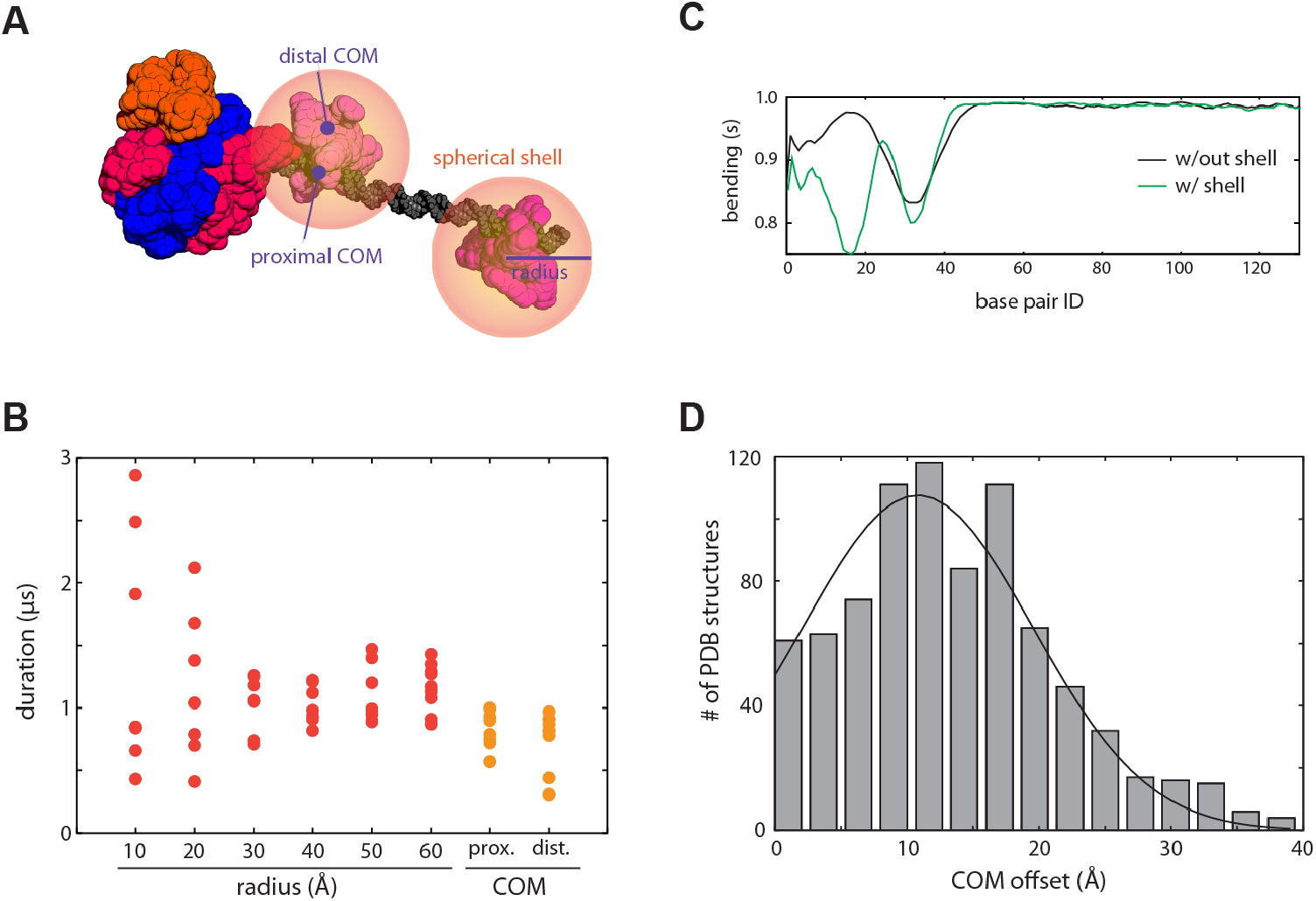
Molecular dynamics simulations of RecBCD collisions with EcoRI molecules surrounded by spherical shells. **(A)** Example of an initial structure used in the molecular dynamics simulations. In the simulations, we varied the radius and center of mass (COM) of the spherical shells to eliminate any protein surfaces or binding geometries that are specific to EcoRI, thereby allowing us to emulate a “generic” DNA binding protein. **(B)** Average duration between the collision of the two proteins molecules and the dissociation of the protein adjacent to RecBCD. **(C)** Representative DNA bending score for proteins encased within a spherical shell; in all cases protein dissociation from the DNA arose from extensive torque–induced DNA bending and distortion of the underlying protein–DNA interface. **(D)** Calculated distances between the protein COM and DNA axis for 933 protein–DNA complexes obtained from the Protein Data Bank (PDB). A Gaussian fit to the data reveals an average COM offset of 10.8 Å.

**Fig. S9.**
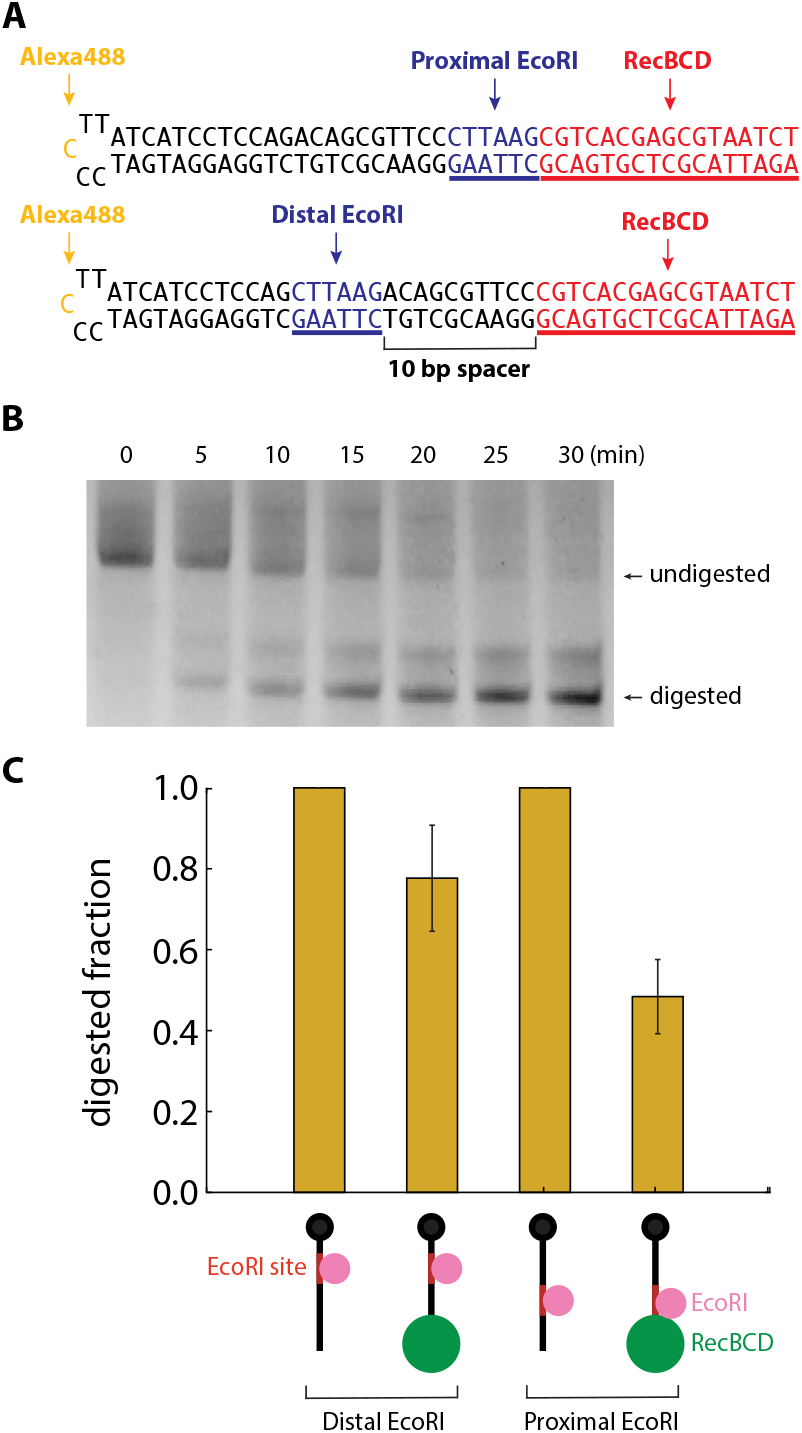
Digestion of DNA oligonucleotides by wild type EcoRI in the presence and absence of RecBCD. **(A)** Schematic of the oligonucleotide substrates containing proximal and distal EcoRI binding sites. The EcoRI binding sites are shown in blue and the RecBCD binding site is shown in red. **(B)** An example denaturing gel showing a time course of EcoRI digestion in the presence of (static) RecBCD. **(C)** The normalized signal intensity of digested fraction at 30 minutes are plotted for each substrate in the presence and absence of RecBCD. Error bars represent standard deviation calculated from three independent experiments. In the cartoon depictions, the DNA is colored black, EcoRI binding sites are red, EcoRI is magneta, and the RecBCD molecules are green.

